# Complete *de novo* assembly and re-annotation of the zebrafish genome

**DOI:** 10.1101/2025.11.17.688901

**Authors:** Javan Okendo, Sergey Koren, Arang Rhie, Aranza Torrado-Tapias, Brandon D. Pickett, Shelise Y. Brooks, Gerard G. Bouffard, Juyun K. Crawford, Christina Sison, Vinita S. Joardar, Terence D. Murphy, Jack A. S. Tierney, Leanne Haggerty, Fergal J. Martin, Catherine Wilson, Angel Amores, John H. Postlethwait, Joy Murphy, Noriyoshi Sakai, Zoltan M. Varga, Adam M. Phillippy, Shawn M. Burgess

## Abstract

The zebrafish (*Danio rerio*) is widely used in vertebrate research, but its reference genome assembly has contained extensively unresolved regions across both euchromatic and heterochromatic compartments. The previous reference genome assembly, GRCz11, consisted of 19,725 contigs assembled into 1,917 scaffolds. Recent advances in both long-read sequencing technologies and genome assembly algorithms have made “complete” genome assemblies possible for the first time. We used homozygous fish from two lab strains, “Tübingen” and ”AB,” for *de novo* genome assemblies. The new assemblies incorporated 7% more genomic sequence than GRCz11 and an additional 130 million bases of previously unassembled sequence. RefSeq annotation incorporating newly generated Iso-Seq cDNA sequences have added notable increases in mRNAs (68%), lncRNAs (47%), and misc_RNAs (1099%). Two assemblies have been elevated to reference genome status (GRCz12tu and GRCz12ab). We generated an additional 40 draft haplotypes to create a zebrafish pangenome resource and demonstrate its utility for variant analysis.

## Introduction

The zebrafish *(Danio rerio),* is the second most commonly used vertebrate model system after mice for studying developmental biology primarily due to their ease of breeding, transparent embryos, and amenability to genetic manipulation and human disease modeling^1–3^. The most recent zebrafish reference genome assembly, GRCz11, was 1.3 Gb and was released in 2017. That assembly consisted of 19,725 contigs assembled into 1,917 scaffolds and was generated with short-read sequencing in combination with a high-density genetic map and Bacterial Artificial Chromosome (BAC) libraries made from the Tübingen strain (TU)^4^. The first zebrafish reference genome project had its origins at the Wellcome Sanger Institute more than two decades ago. This effort faced many technical challenges including extremely high polymorphism rates (greater than 1% between closely related fish), remnants of a whole genome duplication that left approximately 20% of the genes with two copies^5,6^, and a relatively high percentage of repeat sequences and heterochromatin. The zebrafish reference genome has gone through multiple versions, each one representing distinct and significant improvements^7,8^. A particular technical challenge was that the use of BACs caused the repetitive regions of the genomes to be underrepresented as BACs containing a high number of repeats can be unstable and undergo deletions during library preparation or propagation^9^. Due to the many technical limitations, numerous gaps in the GRCz11 assembly remained unresolved and as a result, many of the repetitive and polymorphic regions were either left incomplete or incorrectly assembled.

The current reference genome, GRCz11, is 93% complete and the approximately, 7% of missing genome sequences (containing roughly 100 Mb) are scattered across the genome. These regions include telomeric, sub-telomeric, centromeric and pericentromeric areas, along with ribosomal DNA (rDNA) arrays, all of which are crucial for various biological functions. The recent emergence of new sequencing technologies: PacBio circular consensus sequencing (PacBio HiFi)^10^ and Oxford Nanopore’s Technology (ONT)^11^, now offer a solution to these assembly challenges, enabling the accurate sequencing and placement of the repetitive regions of the genome. These advances in both long-read sequencing technologies and genome assembly algorithms have made complete genome assemblies possible for the first time as has been shown by the recent release of a fully assembled human^12^, mouse^13^, maize^14^, and rat^15^ reference genomes. We assembled three complete haplotype genome sequences from commonly used lab strains, two from homozygous clones generated from the “Tübingen” lab strain one of which is now a reference genome “GRCz12tu.” We also generated a second reference genome for the “AB” lab strain, “GRCz12ab” which was sequenced from the inbred “M-AB” fish^16^. The new assemblies appear gapless, and previous mis-assemblies have been corrected, resulting in the incorporation of an additional 103 million novel base pairs and providing the research community with a highly accurate reference sequences for this important model system.

## Results

### The fully assembled zebrafish genome

The two most commonly used laboratory strains of zebrafish are known as “Tübingen” (TU)^17^, which was the strain used for the original zebrafish reference genome^4^, and “AB”^18^, the strain used in the earliest experiments to establish zebrafish as a vertebrate model system^19^. We used two separate strategies to generate assemblies for each laboratory strain. For the TU assembly, we generated gynogenetic fish using ultra-violet irradiated sperm and a heat shock treatment at mitosis I to generate “doubled haploid,” homozygous diploid fish^19^. For the AB assembly, we used a recently isolated, fully inbred line derived from AB called “Mishima-AB” (M-AB)^16^. Fibroblast cell lines were established from the tail fins of two TU clones and the M-AB line. Genomic DNA isolated from adult tissue were used for PacBio HiFi sequencing and genomic DNA isolated from the fibroblast cell lines was used for Oxford Nanopore sequencing.

We performed whole genome sequencing and *de novo* assembly on three samples; the two assemblies from TU clones obtained from the Zebrafish International Resource Center (ZIRC) and one AB assembly from a M-AB fish obtained from the National Institute of Genetics in Mishima Japan. We selected one of the TU assemblies and the AB assembly and the selected assemblies were elevated as the new reference genomes for *Danio rerio*: “GRCz12tu,” (GenBank accession: GCA_049306965.1) and “GRCz12ab” (GenBank accession: GCA_052040795.1). The third assembly made from the other TU clone was also submitted to GenBank as “NHGRI_Fish11” (GenBank accession: GCA_033170195.3). The assemblies were performed using PacBio circular consensus sequencing (HiFi) and Oxford Nanopore Technology (ONT) ultra-long read sequencing. Genome assembly was conducted using Verkko v. 2.2 with semi-manual corrections where ultra-long (> = 100kb) ONT reads were used for gap patching^20,21^. The GRCz12tu, NHGRI_Fish11, and GRCz12ab assemblies each span approximately 1.4 Gb, an increase of approximately 103 mega bases relative to the GRCz11 assembly (Fig. 1A and Supplementary Table S4). Additionally, we successfully closed all 20,233 gaps that remained unresolved in GRCz11^4^, making all three genome assemblies completely gap-free with all 25 chromosomes now having both full centromeric and telomeric sequences (Fig. 1A and Supplementary Table S5). Additionally, all 968 unplaced contigs and 28 Mb of previously unassembled bases have been fully integrated into the assembly (Supplementary Table S5). Using Merqury^22^, the genomic quality value (QV) was estimated to be 56.9 for GRCz12tu, averaging better than one error in 500,000 bases. The individual chromosomal QV ranged from 53.5 to 62 (Supplementary Table S1). Previous mis-assemblies, primarily because of complex heterochromatic sequences, particularly on chromosome 4, are now fully resolved (Figs. 1C and D). The NHGRI_Fish11 genome assembly (the second TU assembly) achieved a QV of 52 with chromosomal values ranging from 42.64 to 61.29 (Supplementary Table S2). The GRCz12ab genome assembly scaffold achieved a QV of 59.72, approximately 1 error per 1,000,000 bp, with the chromosomal QVs ranging from 53.71 to 63.17 (Supplementary Table S3).

**Fig. 1:**
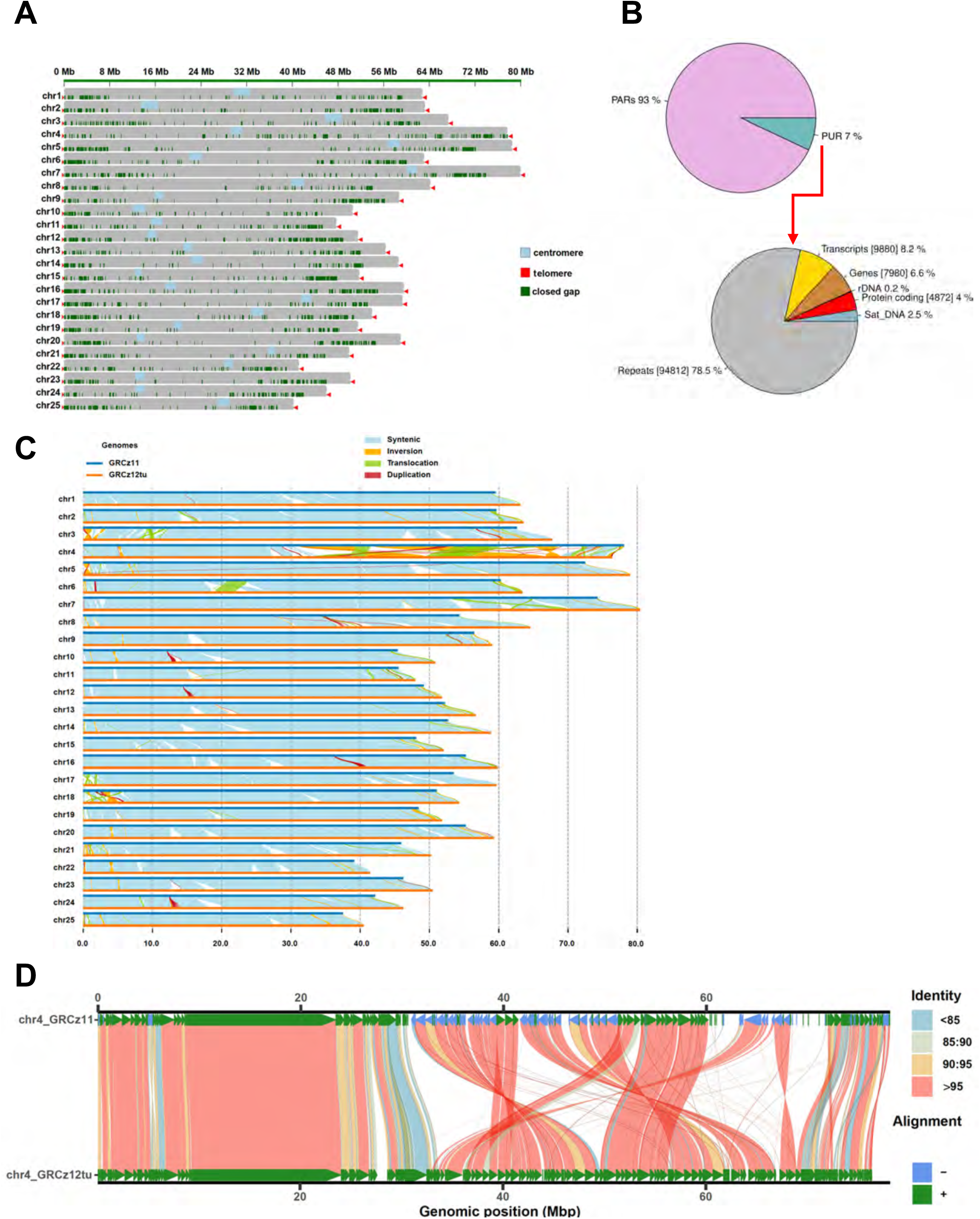
Complete zebrafish genome assembly. (**A**) An ideogram plot showing resolved issues with the GRCz11 assembly. The new components shown on the plot are centromeres (light blue), telomeres (red), and all the closed gaps (green). (**B**) The distribution of genomic elements found in previously unassembled regions (PURs) of the GRCz11. Pie chart showing the proportion of previously assembled regions (PARs) and the previously unassembled regions (PURs). (**C**) Synteny analysis using SyRI^59^ software identified shared chromosomal regions with the greatest percentage increase in chromosomal lengths in the GRCz12tu compared with GRCz11. (**D**) SVbyEye^46^ plot showing the alignment of chromosome 4 from GRCz11 and GRCz12tu genome assemblies. The resolution of inverted segments in chromosome 4 that are now resolved in GRCz12tu are highlighted.

The k-mer distribution plots from GRCz12tu, NHGRI_Fish11, and GRCz12ab assemblies are made up of > 99% of 1-copy k-mers an indication that both assemblies have minimal artifacts from mis-assemblies. The PacBio HiFi and the ONT mapping across both TU haplotypes and the AB haplotype indicate broadly uniform coverage of the reads to the 25 chromosomes in the GRCz12tu, NHGRI_Fish11, and GRCz12ab assemblies (Supplementary Figs. 2, 3 and 4). Some anomalies in ONT coverage are likely due to chromosome number instabilities observed in the cultured fibroblasts (e.g. ONT coverage for chr5 in the GRCz12tu assembly) resulting in some chromosomes being oversampled.

The GRCz12ab genome has a total scaffold length of 1.5 Gb (Supplementary Table S5). In comparing the GRCz12ab with the recently published pseudohaplotype assembly from of an AB strain, DrAB1 (GCA_020184715.1)^23^, DrAB1 had 164 contigs compared to the 25 in GRCz12ab. We also observed an improvement in scaffold contiguity compared to a recently assembled pseudohaplotype generated by the Tree of Life Programme: fDanRer4.1^24^ (Supplementary Table S3). The GRCz12ab and GRCz12tu assemblies are more than 95% syntenic to each other with some instances of significant segmental differences (Supplementary Fig. 17).

The new assemblies show an increase in chromosomal length for 24 chromosomes, all except for chromosome 4, which was reduced in size for all fully sequenced haplotypes compared to GRCz11 (Fig. 1C). However, the extent of this increase is not uniform across chromosomes (Supplementary Table S4). The Chromosome 4 assembly experienced a length reduction of -2.20%, -2.84 and -3.19% in GRCz12tu, GRCz12ab, and NHGRI_Fish11 respectively (Fig. 1C and Supplementary Table S4), while the remaining 24 chromosomes exhibited length increases ranging from 4% to 11.96% (Supplementary Table S4). The decrease in chromosome 4’s length is attributed to the resolution of inverted repeat expansions (caused by mis-assembly) that existed in GRCz11 due to technological constraints^25^. Dramatic changes in the orientation and location of many contigs in chromosome 4 have been identified and corrected (Fig. 1D, Supplementary Fig. 8). Most of the chromosomal length increases in the other chromosomes were concentrated near the sub-telomeric ends. Additionally, we identified many small local differences, such as inversions, translocations, and duplications (Fig. 1C and Supplementary Fig. 1). These presumed structural variations could represent actual strain-specific segmental differences or mis-assemblies (Fig. 1C).

We assessed the conservation of sequence and conserved synteny between both TU haplotypes: GRCz12tu and NHGRI_Fish11 (Supplementary Fig. 1). Synteny analysis revealed a high degree of preservation in genomic order and orientation between the two assemblies (Supplementary Fig. 1). At the chromosomal level, GRCz12tu and NHGRI_Fish11 exhibit substantial structural similarity, although chromosomes 4, 6, and 8 contain large segmental insertions that show significant differences in chromosomal structure even with relatively closely related zebrafish individuals (Supplementary Fig. 1). There were 1,031,073 SNVs between GRCz12tu and NHGRI_Fish11. Because these fish were generated by gynogenesis from the same female fish, this rate of polymorphism is significantly lower than what is seen in natural populations where SNV rates run 1–4% depending on the comparison^26^. In the current GRCz12tu genome assembly, previously assembled regions (PARs) account for 93% of the genome, while the previously unassembled regions (PURs) constitute 7% of the total genome sequence (Fig. 1B). A substantial portion of the PURs consisted of repetitive sequences, which made up 78.5% of these previously unresolved regions (Fig. 1B).

The GRCz12tu assembly exhibits uniform coverage across chromosomes, with a few instances of coverage dips (Supplementary Fig. 2). Further investigations revealed that these dips are in the centromeric regions and locations having functional segmental duplications that are rich in repetitive sequences (Supplementary Fig. 6). In contrast, the NHGRI_Fish11 assembly shows some instances of coverage variation across specific chromosomes. For instance, ONT and PacBio HiFi coverages were similar for chromosomes 1 and 7, but other regions displayed variability (Supplementary Fig. 3). We observed some chromosome number instability in the fibroblast line from NHGRI_FISH11 and the changes in coverage may reflect chromosomal copy number changes in those cultures. In the GRCz12ab assembly, quality ONT read coverage was significantly higher at 184X coverage than that of the PacBio HiFi reads which was at 25X coverage (Supplementary Fig. 4).

The PURs contained a total of 7,980 genes, 4,872 of which were protein coding (Fig. 1B). The PURs also harbored ribosomal DNA (rDNA) arrays and satellite DNAs, both of which have functional roles in genome stability, gene regulation, and chromosome structure^27,28^. The GRCz12ab assembly was compared to both the GRCz12tu assembly and a previously published pseudo-haploid assembly for AB (Dr1AB^23^). The collinearity of GRCz12tu and GRCz12ab was very high while DR1AB, despite being assembled from fish originally derived from the same background as GRCz12ab, aligned more poorly to it, particularly chromosome 4, but also notable differences in chromosomes 3, 5, and 7 (Supplementary Fig. 16).

### Characterization of newly assembled genome components

The new assemblies resolve previously inaccessible genomic regions characterized by low complexity and high repeat content allowing us to characterize regions with extreme levels of heterochromatin. In all three assemblies, we resolved all 25 centromeres as revealed by both CentroMiner, which is part of the quarTeT tool-suite^1^ and StainGlass^29^ software analysis and that gave consistent centromeric repeat coordinates (Figs. 2A, B, C, Supplementary Fig. 5, and Supplementary Table S7). The GRCz12tu centromeric lengths ranged from ∼1 Mb to 2.92 Mb (Supplementary Fig. 10, Supplementary Table S7). Comparing zebrafish with other model systems like mouse ^13^ and rat ^30^, which have telomeric and telocentric centromeres, zebrafish centromeres were all submetacentric (Supplementary Fig. 7) confirming cytogenetics ^31–35^.

**Fig. 2.**
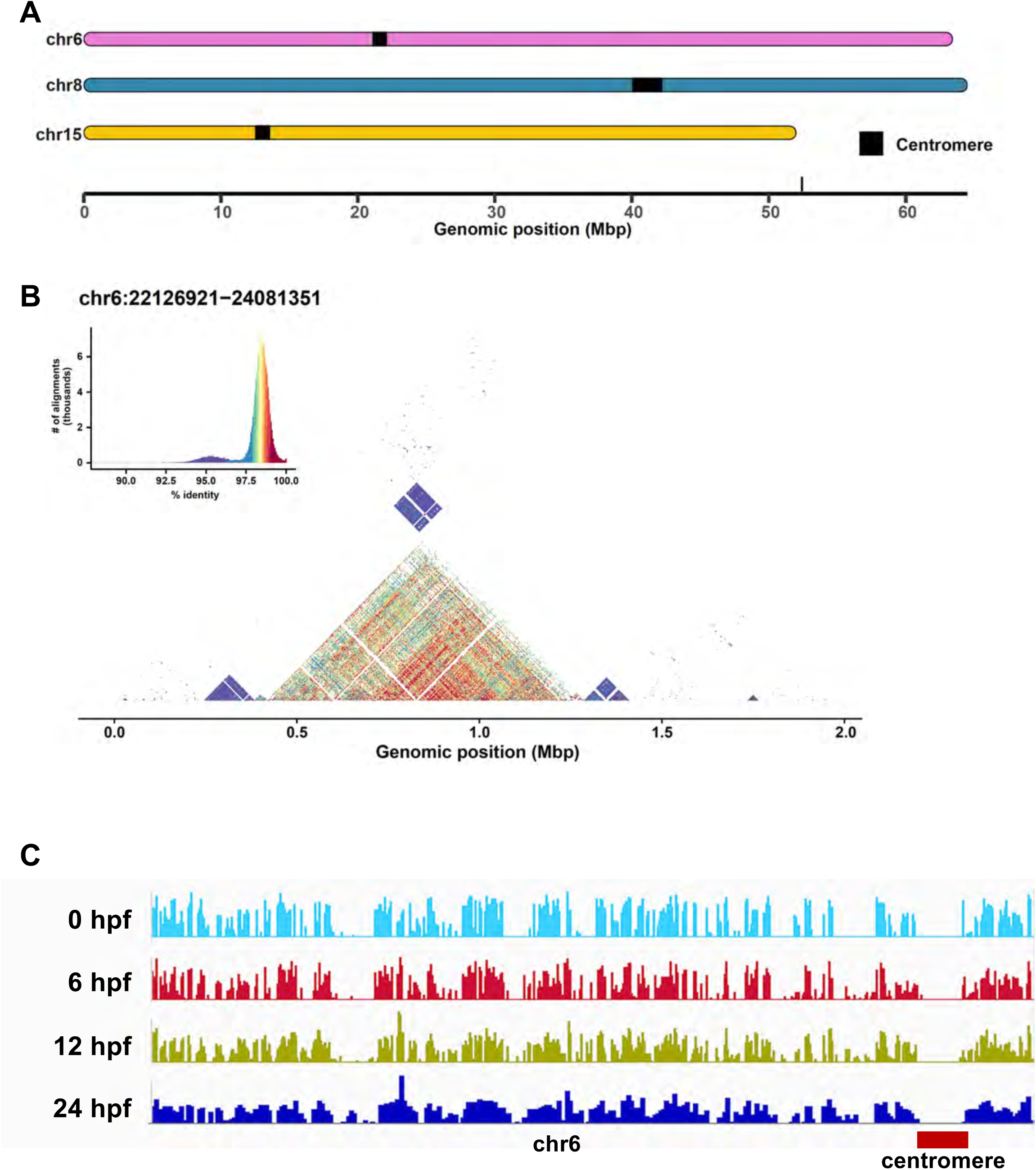
Centromere analysis. GRCz12tu centromere locations as visualized by SVbyEye^60^. (**A**) Centromere chromosomal locations for chromosomes 6, 8, and 15. (**B**) We utilized the StainedGlass ^29^ software to analyze self-to-self similarity within chr6 centromere region to detect centromeric satellite sequences. Dark red shades represent nearly 100% identity, while purple shades indicate similarity levels up to 95%. (**C**) The alignment of Iso-Seq reads from 0,6,12, and 24 hours post-fertilization (hpf) to the short arm of chromosome 6. There is a noticeable lack of transcript alignments spanning the centromeric region. Additional details for other chromosomes are provided in Supplementary Table S7 and Supplementary Fig. 6.

The distribution of centromeric satellite repeats varied among the chromosomes (Fig. 2A, Supplementary Fig. 5). For instance, chromosome 3 exhibited a stable and well-defined arrangement of satellite repeats concentrated in the core centromeric region (Supplementary Fig. 5). In contrast, other chromosomes, such as chromosomes 5, 16, and 21, show evidence of non-satellite repeat incorporation, suggesting a more complex centromeric architecture^36^ (Supplementary Figs. 5 and 6). The variation in alphoid-DNA sequence in our data is consistent with human studies where chromosomal homogenization of the alpha-DNA sequences in the centromere and pericentromeric regions showed significant variation^37^. In humans, the rearrangements in the centromeric cores can cause the insertion of non-satellite DNA sequences as was demonstrated on chromosome 21^37^. We identified the insertion of non-satellite repeats such as long tandem repeats, short interspersed nuclear elements (SINEs), long interspersed nuclear elements (LINEs) and long tandem repeats in all 25 centromeres of the GRCz12tu assembly (Supplementary Fig. 6). The chromosome 1 non-satellite DNA had an unusually large array of LTR/ERV1 sequences in addition to other non-satellite sequences marking it as the most complex centromere (Supplementary Fig. 6).

In GRCz12tu, the total centromere sequence length from the 25 chromosomes was 45.53 Mb while in NHGRI_Fish11 the total centromere length was 48.12 Mb, a difference of 2.59 Mb (Supplementary Table S7). Similar size differences have been seen in human samples^38^. There was notable variation in the centromere length for the same chromosome in different haplotypes. The largest difference variation was observed in chromosome 15 where the two TU haplotypes have a total centromeric length difference of more than 1 Mb and the GRCz12tu and GRCz12ab reference genomes had a 0.5 Mb difference in total centromere length (Supplementary Fig. 10). The presence of satellite DNA sequences on all chromosomes was confirmed through StainedGlass ^29^ analysis, which employs sequence homology and motif recognition to visualize and characterize tandemly repeated DNA regions (Fig. 2B). Unsurprisingly, cDNA sequences were largely unrepresented in the predicted centromeric regions (Fig. 2C).

The paradise fish (*Macropodus opercularis*) has a highly compressed genome of 0.6Gb and its draft assembly^15^ contained five fully sequenced centromeres of the 23 total. We compared the paradise fish centromeric sequences to zebrafish centromeres. The zebrafish centromeres averaged 1.82 Mb, whereas paradise fish centromeres were consistently shorter, all measuring below 0.5 Mb with an average of 0.1 Mb. In zebrafish, the centromeres exhibit a complex architecture with a higher abundance of non-satellite repeat insertions compared to the paradise fish (Supplementary Fig. 12).

We examined the telomeric regions to identify the hexamer repeat sequences characteristic of telomeres in all three assemblies (Supplementary Table S8). We observed variation in chromosomal telomeric sequence lengths with GRCz12tu chromosome 14 having the longest telomeric sequence length of >20kb (Supplementary Table S8). The telomere repeat sequences (TTAGGG)n are the same as has been described for most vertebrates, including humans^39^.

### Structural variations between zebrafish strains

Comparing the GRCz12tu, NHGRI_Fish11, and GRCz12ab genome assemblies revealed key structural variations including deletions, insertions inversions, and transpositions across the genomes (Fig. 3 and Supplementary Fig. 9). These variations are spread across chromosomes, with deletions and insertions being most concentrated in the centromeric and pericentromeric regions of the chromosomes (Supplementary Fig. 9). When comparing NHGRI_Fish11 to GRCz12tu, NHGRI_Fish11 had a total of 606 structural variants of which 597 (98.7%) were insertions and 9 (1.3%) were deletions (Fig. 3 A and Supplementary Table S10) relative to GRCz12tu. The mean insertion length was 6,713 (6.7 kb) and the minimum and the maximum insertion length ranged from 77 bp to 129,745 bp with a median length of 5,290 bp (Table 1, Fig. 3B). The deletions detected in the NHGRI_Fish11 ranged from 55 bp to 4,718 bp with the mean deletion length being 1,174 bp and the median length totaling 236 bp (Table 1). Comparing the GRCz12ab to GRCz12tu, there were a total of 2,943 (99.7%) insertions and 21 (0.3%) deletions, (Fig. 3 A and Supplementary Table S 10). When we compared the GRCz12tu and GRCz12ab genome assemblies, we identified 10kb average insertion sequence lengths and 609 bp for deletions (Table 1 and Fig. 3B). We then set to compare the proportion of structural variants (SV) in the assemblies, and our data shows that insertions were the most abundant SV type in both comparisons at 99.3 % and 98.5 % in GRCz12tu to GRCz12ab and GRCz12tu to NHGRI_Fish11 comparisons respectively.

**Fig. 3:**
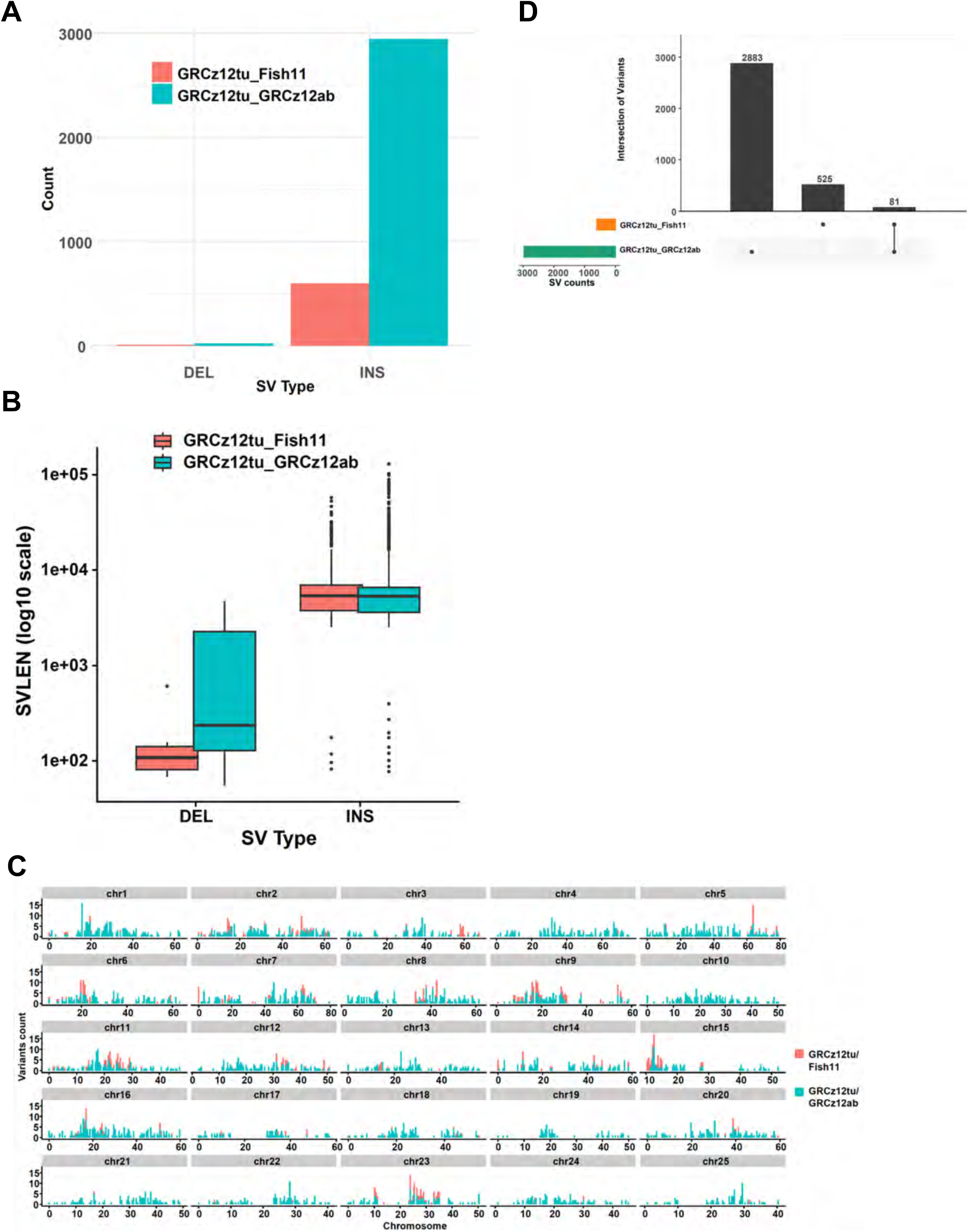
Characterization of structural variants from the three zebrafish assemblies. (**A**) Bar plot showing the total structural variant counts by type (DEL and INS) comparing GRCz12tu to NHGRI_Fish11 (red) and GRCz12tu to GRCz12ab (cyan). GRCz12ab was 1.9 mbp larger than GRCz12tu, reflected in the high number of inserted sequences. (**B**) boxplots highlighting the mean length distribution of deletions (DEL) and Insertions (INS) in the comparisons between the GRCz12tu/GRCz12ab and GRCz12tu/NHGRI_Fish11 genomes. (**C**) The genomic distribution of structural variations for each chromosome comparing GRCz12tu to NHGRI_Fish11 (cyan), and GRCz12tu to GRCz12ab (red). (**D**) UpSet plot showing the proportion of unique and shared SVs in GRCz12tu to GRCz12ab and the GRCz12tu to NHGRI_Fish11 comparisons. Only 81 SV were shared between NHGRI_Fish11 and GRCz12ab emphasizing the extreme variation found in the various sequenced genomes.

**Table 1.**
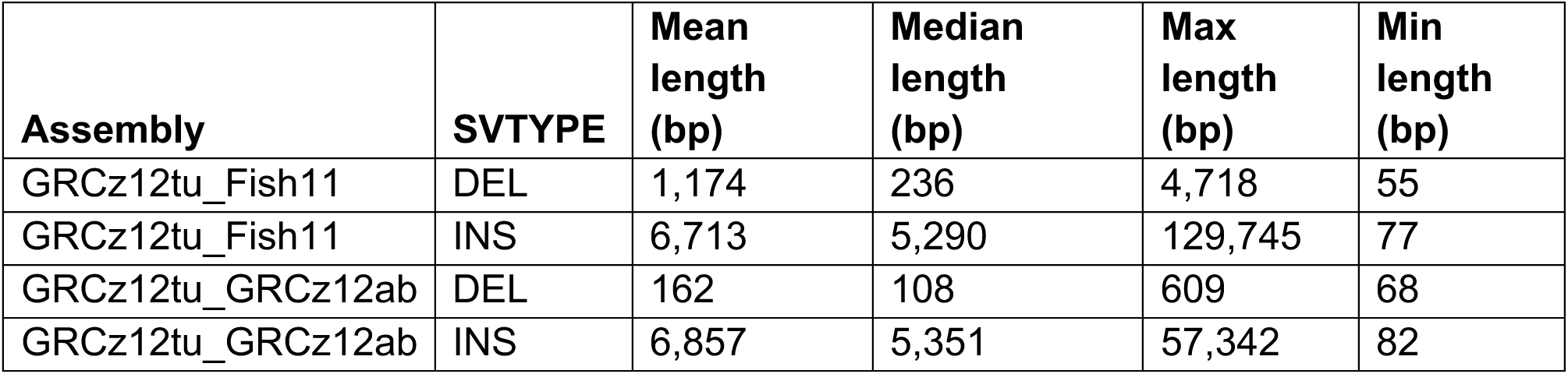
Segment-wise comparisons of GRCz12tu, NHGRI_Fish11, and GRCz12ab genome assemblies highlighting the summary statistics of the insertions and deletions identified when GRCz12tu was compared with NHGRI_Fish11 (another TU haplotype) and the comparison of structural variants arising from the comparison of the GRCz12tu and GRCz12ab genome assemblies. The table represents a detailed analysis of the different genomic variants in different segments of our genome assemblies, offering insights into the structural and sequence-level variations.

| **Assembly** | **SVTYPE** | **Mean length (bp)** | **Median length (bp)** | **Max length (bp)** | **Min length (bp)** |
| --- | --- | --- | --- | --- | --- |
| GRCz12tu_Fish11 | DEL | 1,174 | 236 | 4,718 | 55 |
| GRCz12tu_Fish11 | INS | 6,713 | 5,290 | 129,745 | 77 |
| GRCz12tu_GRCz12ab | DEL | 162 | 108 | 609 | 68 |
| GRCz12tu_GRCz12ab | INS | 6,857 | 5,351 | 57,342 | 82 |

We detected a wide distribution in the frequency of structural variants across the 25 chromosomes between the two, fully assembled TU haplotypes (Fig. 3C). The same trend has been observed in humans where certain genomic locations, specifically those rich in segmental duplications and transposable elements, harbor more structural variation^40^. The highest number of detected variations occurred on chromosomes 2, 9, and 23 (Fig. 3C). Comparing GRCz12tu to GRCz12ab, a different pattern is seen where chromosomes 1, 5, 7, and 16 had the highest number of variant counts. We observed inversions between the GRCz12tu and NHGRI_Fish11 (14 total) and between the GRCz12tu and GRCz12ab genome assemblies (100 total) (Supplementary Table S10). A total of 162 (GRCz12tu and NHGRI_Fish11) and 1,184 instances (GRCz12tu and GRCz12ab) of transpositions were detected, indicating genome rearrangements between haplotypes (Supplementary Table S10). Our data showed 6,084,598 single nucleotide variants (SNVs) between the GRCz12tu and GRCz12ab haplotypes (Supplementary Table S10). Duplications were also detected and were most often located in the repetitive regions of the genome near or in centromeres (Supplementary Table S10). We observed a similarity in the distribution of variants between the assemblies, an indication of shared structural variation hotspots in zebrafish. Specifically, chromosomes 2, 5, 9, 11, 15, and 23 (Fig. 3C) showed higher than average variation. Haplotype-specific enrichment of variants was also observed on chromosomes 10 and 19 (Fig. 3C). Regions of both hotspots and low rates of variation were observed in our data (Fig. 3C). The high degree of structural variation between GRCz12tu and NHGRI_Fish11 shows that even within zebrafish lab strains, there is significant variation that could impact measures of gene expression and analysis of phenotypes. Merging the structural variant analysis between GRCz12ab and GRCz12tu, we observed a total of 2,883 variants comparing GRCz12tu to GRCz12ab, and a total of 525 SVs that were present comparing GRCz12tu to Fish11. Only 81 of the detected SVs were shared between NHGRI_Fish11 and GRCz12ab (Fig. 3D).

**Table 1**. Segment-wise comparisons of GRCz12tu, NHGRI_Fish11, and GRCz12ab genome assemblies highlighting the summary statistics of the insertions and deletions identified when GRCz12tu was compared with NHGRI_Fish11 (another TU haplotype) and the comparison of structural variants arising from the comparison of the GRCz12tu and GRCz12ab genome assemblies. The table represents a detailed analysis of the different genomic variants in different segments of our genome assemblies, offering insights into the structural and sequence-level variations.

### NCBI RefSeq gene annotation

Gene annotations for GRCz12tu were generated using NCBI’s Eukaryotic Genome Annotation Pipeline^41^ with a combination of curated data and computational models based on extensive, publicly available, short-read, long-read, and CAGE RNA-seq data. The annotation set, referred to as GCF_049306965.1-RS_2025_04, includes 27,039 protein-coding, 22,113 non-coding, and 406 pseudogenes (Table 2). A total of 99.5% of transcripts are fully supported by aligned transcript or protein sequences, including 73% of Gnomon transcript models with 5’ ends based on CAGE data and 74% with 3’ ends based on polyadenylated transcript evidence. BUSCO analysis of the annotation, using one longest protein per gene, indicates that the annotation is 99.0% complete using the actinopterygii_odb10 dataset with 3,640 single-copy models.

**Table 2.** Comparison of National Center for Biotechnology and Information Reference Sequence (NCBI RefSeq) annotations for GRCz11* and GRCz12tu. GRCz11**\*** statistics reported for the primary assembly only, not including alternate loci scaffolds representing sequence from other strains.

| Feature |  | GRCz11* | GRCz12tu |
| --- | --- | --- | --- |
| Genes and Pseudogene |  | 45,520 | 49,597 |
|  | Protein-coding | 26, 862 | 27,039 |
|  | Noncoding | 18,187 | 22,113 |
|  | Pseudogenes | 431 | 406 |
|  | Genes with variants | 11,172 | 18,413 |
| mRNAs |  | 49,400 | 89,815 |
|  | Known RefSeq (NM_) | 16,624 | 16,543 |
|  | Model RefSeq (XM_) | 32,776 | 73,272 |
|  | Partial | 1,244 | 179 |
|  | Models with filled gap(s) | 605 | 0 |
|  | Known RefSeq split across multiple locations | 67 | 2 |
| Noncoding RNAs |  | 21,501 | 41,222 |
|  | Known RefSeq (NM_) | 631 | 623 |
|  | Model RefSeq (XM_) | 12,352 | 31,872 |
| CDSs |  | 49,428 | 89,857 |
|  | Known RefSeq (NM_) | 16,622 | 16,555 |
|  | Model RefSeq (XM_) | 32,776 | 73,272 |
|  | With > 5% *ab initio* | 770 | 178 |
|  | Partial | 1,135 | 77 |
|  | Know RefSeq with CDS coverage < 90% | 397 | 74 |
|  | With major correction(s) | 724 | 211 |
| All transcripts |  | 70,901 | 131,037 |
|  | mRNA | 49,400 | 89,815 |
|  | misc_RNA | 983 | 13,617 |
|  | miRNA | 557 | 559 |
|  | tRNA | 8,312 | 8,518 |
|  | lncRNA | 7,264 | 12,861 |
|  | snoRNA | 246 | 260 |
|  | snRNA | 1081 | 1,283 |
|  | Antisense RNA | 2 | 2 |
|  | rRNA | 3,047 | 4,113 |
|  | Rnase MRP RNA | 1 | 1 |

Compared to the most recent RefSeq annotation of GRCz11 from August 2024 (GCF_000002035.6-RS_2024_08), the GRCz12tu annotation has a 0.6% increase in protein-coding genes and a 22% increase in non-coding genes (Table 2). A sub-telomeric rDNA array on chr4 expanded from one to eighteen 45S rDNA units, and a second array on chr5 expanded from two partial units to ten units. A complex region on chr8 composed predominantly of DWNN domain-containing genes related to the N-terminus of RBBP6 expanded from 600 kb (NC_007119.7: 34,434,345-35,034,401) to 3.7 Mbp (NC_133183.1: 36,572,689-40,237,669), increasing the number of RBBP6-like genes from 18 to 208 copies.

In addition to the improved assembly, the GRCz12tu annotation utilized a large Iso-Seq dataset including cDNAs from 8 organs (brain, eye, muscle, inner ear, testis, ovary, liver, and kidney) produced as part of this project (SRA project SRP568323), which helped increase representation of lncRNA genes (+50%) and expanded representation of alternatively spliced mRNAs by 68%. Some genes exhibit extensive alternative splicing, with 262 genes exceeding a 50 models/gene threshold imposed on the final RefSeq annotation dataset. The most extensive alternative splicing observed was for *rims2b* (GeneID:563393, ZFIN: ZDB-GENE-060503-893), a regulator of synaptic membrane exocytosis, with 48 exons and 970 predicted transcript isoforms in the initial Gnomon model dataset.

The GRCz12tu annotation shows substantial improvements in gene model quality resulting from closing assembly gaps and increasing sequence quality compared to GRCz11. There are fewer partial mRNAs (87% decrease), which include both known RefSeq transcripts with partial alignments to the genome and model RefSeqs where transcript sequence data was previously used to compensate for exons missing in assembly gaps. We identified 65 RefSeq transcripts that were aligned across multiple disjointed locations in GRCz11 (*e.g.* an inversion or other assembly error) and are now resolved. Both known and model CDSs with major corrections were identified, such as frameshifting indels or internal stop codons, which are substantially reduced (72% decrease). For example, *cep128* (GeneID: 100037385, ZFIN:ZDB-GENE-070410-34), a coiled-coil domain containing ATPase involved in centrosome function^42^, was affected by a 30 kb inversion in GRCz11 that is now resolved (Fig. 4D), and *zgc:76872* (GeneID:403007, ZFIN:ZDB-GENE-040426-1794), an ortholog of the human zinc finger and BTB domain containing protein ZBTB45, was disrupted by a 2 kb gap in GRCz11 that affected four exons and 70% of the protein length (Fig. 4E). The remaining known RefSeq annotations with multiple alignments (2) or partial coverage (74), or both known and model RefSeq annotations with major corrections (211), likely include some annotation errors due to poor predictions or strain variation, which will require curation review. Overall, we estimate that 6% of protein-coding genes benefit from the improvement in assembly quality in GRCz12tu.

**Fig. 4.**
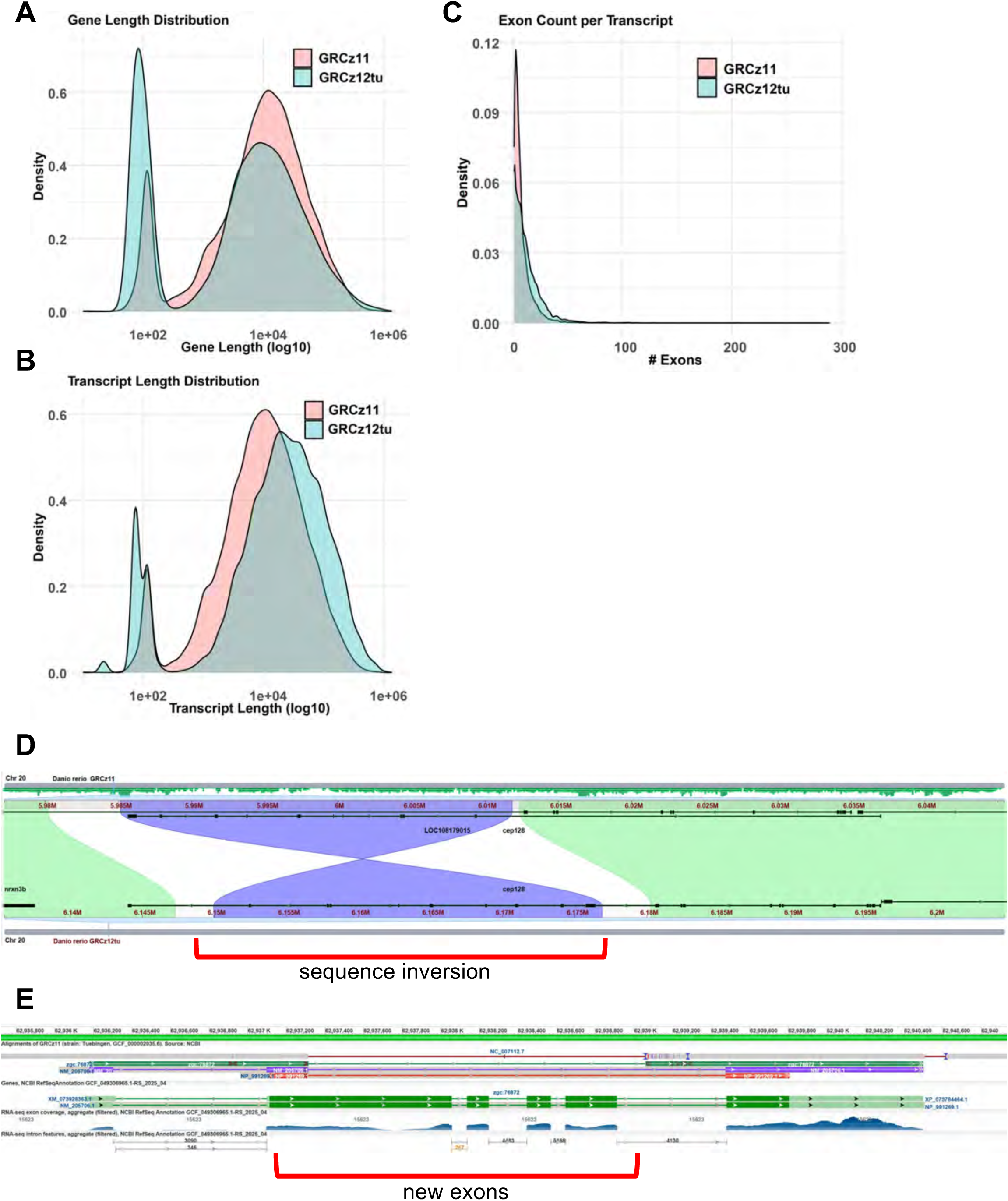
Comparison of gene annotation features between zebrafish genome assemblies GRCz11 and GRCz12tu. Density plot illustrates the distribution of (**A)** gene length, (**B**) transcript length, and (**C**) exon counts per transcript. Both assemblies show similar overall patterns, however, GRCz12tu exhibits modest enrichment for longer gene and transcript lengths (**A** and **C**), suggesting improvements in gene model resolution. The exon counts remain consistent across the assemblies while GRCz12tu has slightly more transcript numbers compared to the GRCz11 assembly. (**D**) The region around *cep128* shows a reversal of the intronic signal eliminating an artifactual transcript and defragmenting *cep128*. (**E**) *zgc:76872* shows an extension of one exon and the addition of three more previously misassembled exons.

Previously unincorporated contigs from GRCz11 have been successfully integrated and the newly resolved sequences within these regions have also been fully annotated.

The gene and transcript length distributions are bimodal in the current reference as well as in the new assembly, with the new assembly showing more transcripts of shorter length and fewer transcripts with longer sequences (Figs. 4A and B). In terms of the number of exons per gene, both GRC11 and GRCz12tu (RefSeq) had the most representation at one exon, indicating a preponderance of single exon transcripts in GRCz11 and GRCz12tu assemblies and a rapid drop-off in exon counts after five exons (Fig. 4C). Many missing exons from the GRCz11 annotation were corrected and included (examples in supplementary Figs. 15A, B and C).

**Table 2**. Comparison of National Center for Biotechnology and Information Reference Sequence (NCBI RefSeq) annotations for GRCz11* and GRCz12tu. GRCz11**\*** statistics reported for the primary assembly only, not including alternate loci scaffolds representing sequence from other strains.

### Segmental duplications and rRNA copy numbers

We quantified segmental duplications (SDs) in GRCz11, GRCz12tu, NHGRI_Fish11, and GRCz12ab haplotypes by identifying SDs with length equal or greater than 20kb that are at least 500kb apart ^43–45^ (Supplementary Table S9). The total number of SDs that were > = 20kb in length and separated by > = 500 kb in GRCz11 was 7,467. In GRCz12tu there were 18,822 SDs, NHGRI_Fish11 had 18,311, and GRCz12ab contained 18,997 (Supplementary Table S9). Chromosome 4 had the most presumed functional SDs followed by chromosome 8 (Fig. 5A). The novel SDs were mainly found in the centromeric, pericentromeric and telomeric regions of the genome (Fig. 5A), the remaining SDs were distributed across the genome. Our data show that the presumed functional SDs were in the same locations as gaps (Fig. 5A), a demonstration that these regions were the primary reason the gaps could not be resolved by the older sequencing technologies. We then quantified and compared the rRNA gene copy number from the GRCz11 and GRCz12tu assemblies. There was an increase of 5S ribosomal rRNA gene copy number in GRCz12tu compared to GRCz11 from 9,100 to 12,092, a 32.88% increase (Fig. 5B, Supplementary Fig. 11). There was an increase in the 5.8S ribosomal RNA sub-unit tandem repeats from 36 in GRCz11 to 120 in GRCz12tu (Fig. 5B). The 18S and 28S ribosomal RNA sub-units also registered a significant increase in tandem repeat numbers when we compared the GRCz11 and GRCz12tu assemblies (Fig. 5B).

**Fig. 5.**
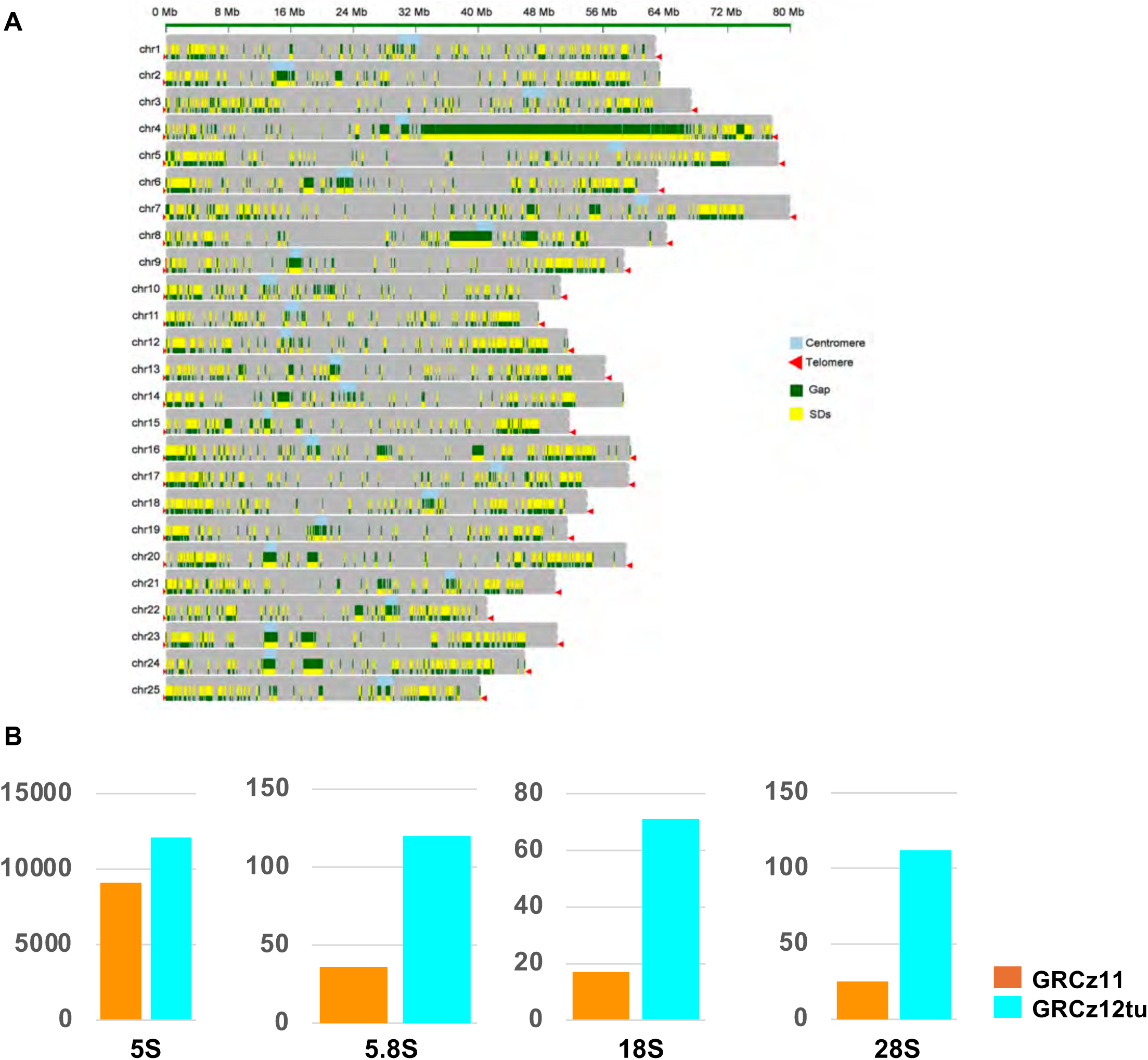
Characterization of segmental duplications throughout the genome. (**A**) Visualization of the GRCz12tu assembly illustrating segmental duplication locations across all the 25 chromosomes. Each horizontal bar represents a single chromosome with segmental duplications plotted along its length. The regions showed include telomeres (**red**), segmental duplication (**green**), and centromeres (**cyan**). Segmental duplications are enriched in the centromeric regions and is the major feature of the long arm of chr4. (**B**) Co-localization of segmental duplications (SDs) with assembly gaps in GRCz12tu assembly. Yellow bars represent SDs, green marks denote assembly gaps, red arrows represent telomeric ends, and light blue represents the centromeric regions. (**C**) Bar plots showing the gene copy numbers of rRNA in the GRCz11 and GRCz12tu assemblies respectively. Note the scale on the Y-axis varies for each rRNA type.

### Zebrafish pangenome construction and comparison of MHC genes

We constructed pangenome graph from 44 haplotypes: our three complete assemblies, fDanRer4.1 (EBI), and draft genomes from 20, wild-caught, normally heterozygous fish using Minigraph-Cactus^46^ and visualized the pangenome segment blocks using ODGI^47^ (Supplementary Fig. 13). The wild-caught fish were captured from two different geographic locations in India. Both wild-caught and laboratory zebrafish strains were included to capture variation across zebrafish populations. The pangenome graph comprised a total node length of 4,347,506,932 base pairs connected with 255,658,588 edges and included 598 distinct paths. Principal component analysis (PCA) based on variants from a zebrafish pangenome Variant Call Format (VCF) reveals a clear stratification among the strains we compared, clustering them into three distinct groups (Fig 6A and Supplementary Fig. 14). Not surprisingly, phylogenetic clustering using the pangenome VCF reveals a distinct separation between either laboratory strain (AB or TU), and wild-caught fish from different locations (CB and EKK) (Fig. 6B). The substantial genetic divergence between wild-caught and laboratory strains of zebrafish is consistent with previous findings^48^ and suggests raising fish in captivity selects for some genetic characteristics regardless of initial origin.

**Fig. 6.**
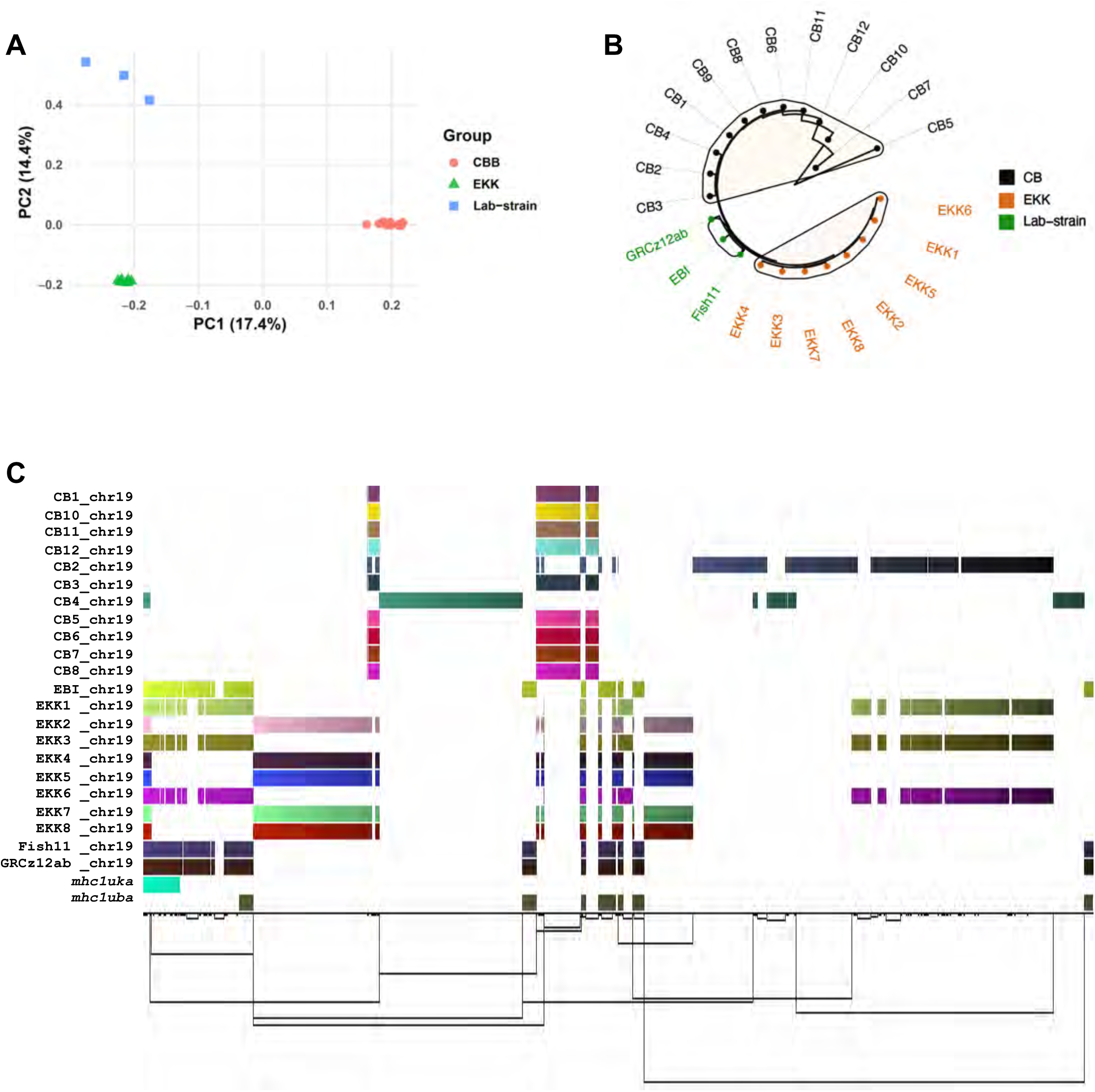
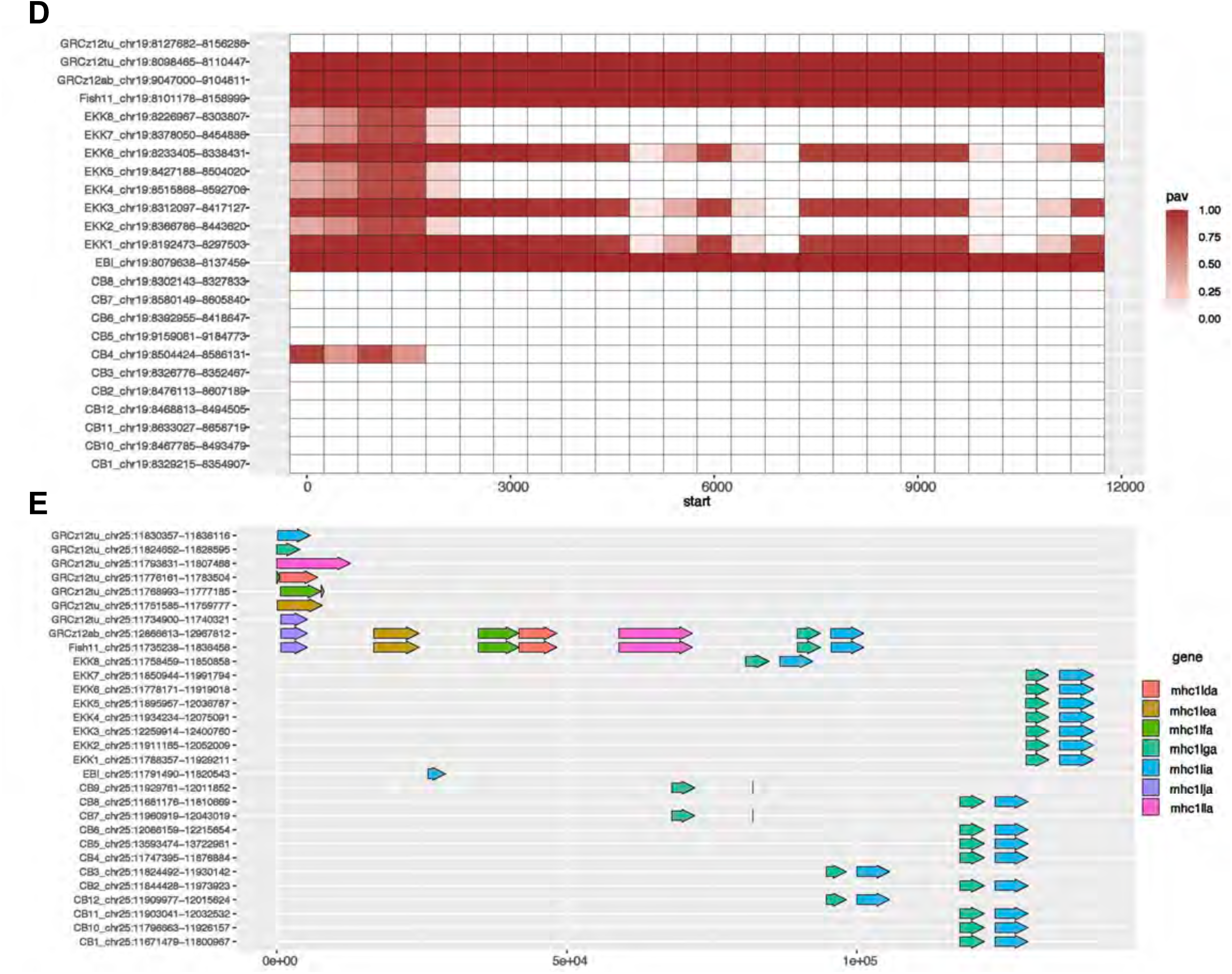
Genomic diversity of wild-caught and laboratory zebrafish strains. (**A**) Principal Component Analysis (PCA) plot displaying the genetic variation among the zebrafish strains we compared using variants obtained from the pangenome multi-sample VCF file reveals substantial genetic differences where each point represents the individual strain. (**B**) Hierarchical clustering using hamming genetic distance further supports the divergence between the wild and laboratory strains, with clear clustering into major clades (**C**) The graphical representation of zebrafish pangenome focused on the mhc1uka gene locus on chromosome 19, reconstructed using sequences from multiple haplotypes demonstrating extensive haplotype diversity. The link at the bottom of the figure shows the presence of structural variation between the haplotypes. Each row in the figure represents the haplotype-specific genome assembly from individual zebrafish, and each color block represents a segment traversed by the haplotype within the pangenome graph. The illustration shows the presence or absence of a particular segment across haplotypes. (**D**) Presence/absence variants visualization showing the genetic divergence and diversity of mhc1uka gene in different zebrafish strains. (**E)** Untangled pangenome graph at a region of chromosome 25 associated with seven different MHC class I genes, showing the arrangements of MHC class I genes arrangements along chromosome 25 of laboratory strains and wild-caught strains.

We then extracted the *mhc1uka* and *mhc1uba* gene loci sub-graph and observed a total graph length of 369,966 base pairs, 12,931 nodes, 17,398 edges, and 24 distinct paths (Fig. 6C). We observed strain-specific path divergence where laboratory strains shared contiguous and non-contiguous alignment blocks that align well with both *mhc1uka* and *mhc1uba* gene annotations (Fig. 6C). In contrast, the wild-caught fish do not traverse the same paths as the GRCz12ab, EBI, Fish11, and GRCz12tu strains through the *mhc1uba* gene locus (Fig. 6C). The wild-caught fish do align well with the *mhc1uka* gene locus (Fig. 6C). The *mhc1uba* is either absent or highly divergent in wild-caught strains while the *mhc1uka* gene appears to be conserved well across laboratory and wild-caught strains but with varied gene copy number (Fig. 6C).

The pangenome construction revealed a presence of variation and structural polymorphism within the *mhc1uka* and *mhc1uba* gene loci across zebrafish strains which was obscured by use of a single strain genome reference for zebrafish^49^ (Fig. 6D). There were distinct genetic presence/absence patterns on chromosome 19. Our data showed that *mhc1uka* gene was variably retained across individual zebrafish, with some fish showing full, partial, or complete absence of sequence regions (Fig. 6C). Some haplotypes shared large contiguous blocks of the presumed conserved sequences, while others exhibited a significant divergence, with alternative segment arrangements, or deletions (Figs. 6C and D). The proximal and distal regions of the *mhc1uka* gene locus exhibited extensive conservation of the segmental structures implying reduced structural diversification, whereas the central region of the gene showed high structural diversity (Fig. 6C). Fish GRCz12tu, GRCz12ab, EBI, and Fish11 shared a unique segmental combination not found in the other haplotypes, an indication of lineage specific variation or loss of heterozygosity in laboratory strains. These difference in the absence/presence of segmental blocks from different zebrafish haplotypes confirmed the idea that zebrafish MHC class I genes, specifically the *mhc1uka* gene were highly polymorphic, with substantial copy number and structural variation between zebrafish haplotypes. The pangenome graph suggests that zebrafish harbor significant *mhc1uka* gene haplotype complexity consistent with what has been observed in humans ^47,50^. The GRCz12tu assembly has all seven classical MHC class I genes arranged in a non-contiguous manner (Fig. 6E). Chromosome 25 had a simplified gene structure in wild-caught strains that encoded the *mhc1lia* gene in tandem copies and the rest of the MHC genes were missing in the sequenced wild-strains (Fig. 6E). The analysis of the *mhc1uka* and *mhc1uba* gene diversity showed structural diversity and copy number variation in zebrafish mirroring what has been observed elsewhere^51^. For instance, the large-scale variation in the human leucocyte antigen (HLA) gene content was only uncovered with the advent of pangenome frameworks that showed that HLA haplotypes differed by several megabases between individual humans^50^.

## Discussion

We report three complete *Danio rerio* “telomere-to-telomere” genome assemblies that bridge previously inaccessible genomic regions and close over twenty-thousand gaps in the current reference assembly. These assemblies include resolution of all telomeres and centromeres, the closure of all the assembly gaps in GRCz11 and the incorporation of all the unplaced contigs into chromosomes. We have identified structural duplications and significant variation both at the nucleotide and structural level between the complete assemblies. Our assembly work confirmed that the unresolved or mis-assembled regions in GRCz11 were mainly composed of the repetitive DNA sequences including centromeric and telomeric sequences which were not resolved using the shorter sequence read technologies. Using pangenome analysis, we demonstrated the variation of *mhc1uka* and *mhc1uba* gene copy numbers in wild-caught and laboratory strains of zebrafish. The large degree of variation almost certainly has a major effect on gene expression, and a broader characterization of that variation will give researchers the tools to study this important aspect of gene regulation in greater detail. The complete telomere-to-telomere genome assemblies will provide an essential foundation for the research community and an opportunity to study the evolutionary details of complex genomic structure creation and function in fish species. The correction of the assembly and annotation of 6% of the zebrafish genes substantially improves the accuracy of the systematic functional annotation of the genome (*e.g*. DANIO-CODE^52^) as well as the comparative functionality of the human and zebrafish genomes, which will improve the utility of zebrafish models for human diseases.

## Resource availability

### Lead contact

For additional information, resources, or reagents, requests should be directed to the lead contact.

### Materials availability

This study did not generate any new unique reagents.

### Data and code availability

This whole Genome Shotgun project for GRCz12tu has been deposited at DDB/ENA/GenBank under the accession JBMGRA000000000. The version described in this paper is version JBMGRA010000000. The PacBio HiFi data can be accessed from the SRA database with the accession ID SRR30635750 while the long read ONT accession numbers are: SRR30635752, SRR30635751, SRR30635749, SRR30635748, SRR30635747, SRR30635746, SRR30635745, SRR30635744, and SRR30635742. For the NHGRI_Fish11, the genome assembly GenBank identifier is GCA_033170195.3, and both ONT and PacBio HiFi reads are available on SRA with the BioProject ID PRJNA1029986. The Iso-Seq RNA-seq reads are available on SRA with Bioproject ID PRJNA1232602. The GRCz12ab PacBio HiFi and ONT raw read sequence data are available on SRA under BioProject ID PRJNA1299309. The GRCz12ab Whole Genome Shotgun project has been deposited at DDBJ/ENA/GenBank under the accession JBQAYU000000000. The version described in this paper is version JBQAYU010000000. The software used in the analysis has been referenced clearly in the text. The raw PacBio HiFi data used to build zebrafish pangenome is available under BioProject ID PRJNA1330284.

## Acknowledgments

This research was supported [in part] by the Intramural Research Program of the National Institutes of Health (NIH). The contributions of the NIH author(s) are considered Works of the United States Government. The findings and conclusions presented in this paper are those of the author(s) and do not necessarily reflect the views of the NIH or the U.S. Department of Health and Human Services. SMB was supported by the National Human Genome Research Institute (ZIAHG000183-24). The work of T.D.M and V.S.J. was supported by the National Center for Biotechnology Information of the National Library of Medicine (NLM), National Institutes of Health. J.A.S.T, LH and F.J.M were supported by funding from Wellcome Trust [WT222155/Z/20/Z] and EMBL core funds.

## Author contributions

J.O. performed the statistical and computational analyses and wrote the first draft of the manuscript. A.T.T. generated cell lines, conducted experiments and purified genomic DNA. S.K., A.R., B.D.P., did additional statistical and computational analysis, S.Y.B., G.G.B., J.K.C. did both PacBio and ONT sequencing at the NIH Intramural Sequencing Center (NISC), C.S., C.W., A.A., J.H.P. provided the 20 wild-caught fish used to build pangenome graph, J.A.S.T., L.H., and F.J.M. performed the Ensembl annotation. V.S.J and T.D.M provided the RefSeq annotation, J.M., Z.M.V. generated the double-haploid fish, N.S. bred the original M-AB line and generated the associated fibroblast cell line, A.M.P., S.M.B. were responsible for initial experimental design, supervision of data analysis and for the final manuscript.

## STAR Methods

### Zebrafish husbandry

Wild-type AB (ZL1; stock# 12093), TU (ZL57; stock# zs119000–1), and *mitfa^b^*^692^*^/b^*^692^ mutants (ZL49; “*nacre*,” stock# zs12089.01) were maintained at the Zebrafish International Resource Center (ZIRC) under standard conditions on a 14:10 hour light: dark cycle, as previously described^53^. Recirculating aquaculture systems employed coarse and fine mechanical filtration, biological nitrification, and UV sterilization. Fish were maintained at a stocking density of 5 - 7 fish/L, with tank water exchanged approximately 4 times per hour. Environmental parameters were continuously monitored and remained within the following ranges: temperature, 28.5 ± 1°C; pH, 7.4 ± 0.4; conductivity, 450–500 µS/cm. Water hardness (dGH 5–6) and nitrogenous waste (NH₃/NH₄⁺ ≪ 0 mg/L; NO₂⁻ < 0 mg/L; NO₃⁻ < 20 mg/L) were tested weekly using colorimetric assays. No significant deviations from these parameters were recorded during the animals’ lifespan. Larvae were fed *paramecia* and *artemia*, transitioning to dry feed at the juvenile stage^53^; adults received a mixture of dry feeds as described^53^. All procedures were approved by the University of Oregon IACUC (AUP-18-05), and routine health monitoring was conducted according to established protocols^54,55^.

### UV inactivation of sperm

AB sperm was collected into E400 extender^56^, spread on ice-chilled concave glass dishes, and irradiated using a Phillips TUV 15W/G15 T8 germicidal lamp (30-min warm-up, 254 nm, at 34 cm distance). UV sperm DNA inactivation was calibrated using five exposure durations (100 - 140 s in 10 s increments), with four TU egg batches per timepoint (mean batch size 274 ± 183 s.d.; total 4,943 viable embryos; two low-count batches excluded). A follow-up experiment tested three additional exposure durations (140-160s) with five distinct batches of TU eggs each (total 3,022 embryos; mean batch size 233 ± 143 s.d.). A 160 s exposure was determined sufficient to ensure complete sperm DNA inactivation without damaging sperm cells.

### In vitro fertilization and heat shock

Artificial diploid embryos were produced by fertilizing TU oocytes with UV-inactivated AB sperm, followed by a timed heat shock to suppress the first mitotic division^19,57^. To this end, ten AB (ZL1) males were anesthetized (MS-222, 168 mg/L) and gently stripped for sperm, which was UV-inactivated (160 s, 34 cm) and stored in E400 on ice. AB, TU, and *mitfa^b^*^692^*^/b^*^692^ (ZL49) females were similarly anesthetized and gently stripped for eggs. Haploid development was initiated by mixing oocytes with UV-inactivated AB sperm in a 1:1 mixture of system water and SS300. For diploid induction, embryos were quickly transferred to a 41°C water bath for 2 min at 13 min post-fertilization, then immediately returned to 28.5°C for recovery. Embryos (heat-shocked and untreated controls) were monitored for morphology, pigmentation, and viability at multiple timepoints until 3 - 6 days post fertilization (dpf), when haploid embryos typically perish (Fig. 1).

### Experimental groups and outcomes

Haploid development was induced in TU eggs (batch #4) using UV-inactivated AB sperm. The clutch was divided equally: one half was subjected to heat shock (41°C, 2 min, at 13 min post-fertilization), while the other half remained at 28.5°C (haploid control). UV-inactivated AB sperm was also used to fertilize *mitfa^b^*^692^*^/b^*^692^ (ZL49) eggs, without heat shock (additional pigment control). Fertility, survival, pigmentation, and haploid/diploid morphology were assessed at 4, 24, 48, 96, and 120 hpf. *mitfa* mutant haploids displayed typical haploid features^57^, lacked wild-type pigmentation (Fig. 1A), and expired between 3 - 5 dpf, confirming complete inactivation of the sperm genome. Forty-two TU HS-diploid embryos were reared in the ZIRC nursery^53^ and shipped as young adults to NHGRI for genomic sequencing.

### Cell culture protocol

The primary cell culture was established by zebrafish caudal fin clippings. In brief, the fish were anesthetized using 150mg/L Tricaine. The fin pieces were washed in ethanol, followed by sterile 1X PBS, and subsequently placed in a sanitization solution for transportation from the fish facility to the laboratory. Under the hood, the fins were further cleaned using pre-warmed Leibovitz’s L-15 medium (Gibco Catalog #11415064). Fins were placed in a 1.5ml microtube, 750ul of Trypsin-EDTA 0.05% (Gibco Catalog # 25300-054) was added and incubated, stirring at 300 rpm at 30°C for 30 minutes. After the incubation was completed, the fin pieces were dissected into smaller fragments and placed in a 12-well cell culture plate in L-15 media supplemented with 20% fetal bovine serum, 10µg/ml bovine insulin (Cell Applications Cat#128-100), 0.12mg/ml kanamycin sulfate-100x (Thermo Fisher Cat# 15160054), 2.5µg/ml amphotericin B (ThermoFisher Cat# 15290026). Twenty-four hours later, the fins were washed with 1X PBS and treated with Sigma Accutase cell detachment solution (Catalog # SCR005) for 30 minutes. After detaching the cells, growth media was added, mixed, and placed in a collagen I-coated 6-well plate (Stem Cell Technologies Catalog #100-0362). The growth media was changed every other day, and cells were sub-cultured when they reached approximately 70-80% confluency.

### High Molecular Weight (HMW) DNA Cells

Adherent cells were detached when they reached 70-80% confluency using Trypsin-EDTA (0.05%) (Gibco Catalog #25300054) as a dissociation agent. Cells were resuspended in a growth medium and transferred to a 15ml tube. Cell suspension was centrifuged for 10 minutes at 200g. After centrifugation, the media was removed, and the cell pellet was flash-frozen in dry ice.

For PacBio sequencing, High Molecular Weight DNA (HMW DNA) was extracted from adult tissue using New England Biolabs (NEB) Monarch® HMW DNA Extraction Kit for Tissue (Catalog # T3060L) following manufacturer’s instructions. For Oxford Nanopore sequencing, genomic DNA was extracted from cultured fibroblasts using Oxford Nanopore Ultra-Long DNA Sequencing Kit V14 (SQK-ULK114) following manufacturer’s instructions.

### Verkko assembler

We performed hybrid assembly with Verkko assembler (version 2.2)^20^, using 25X PacBio HiFi and 60-300x ONT ultra-long sequencing data (reads lengths > = 100kb), producing the final assembly of size 1.4 Gb. More than 50% of the chromosomes were telomere-to-telomere as they were resolved in during the first Verkko run. The remaining chromosomes had complex tangles which were then resolved using the ONT reads of more than 100kb. A semi-manual repeat resolution strategy was employed to further resolve the tangles. The re-alignment of the ultra-long ONT reads was done using GraphAligner v1.0.17^58^, the resultant alignment graph was then used to identify the correct traversals. The reads traversing the correct paths were extracted and supplied to Verkko for gap patching. Comprehensive description of sample preparation, library preparation and sequencing, data processing workflow, and bioinformatic data analysis can be found in the supplementary file section.

### RefSeq Annotation

Annotation of the GRCz12tu assembly was generated for NCBI’s RefSeq dataset using NCBI’s Eukaryotic Genome Annotation Pipeline^41^. The annotation, referred to as GCF_049306965.1-RS_2025_04, includes gene models from curated and computational sources for protein-coding and non-coding genes and pseudogenes, and is available from NCBI’s genome FTP site and web resources.

More than half of protein-coding genes (15,469, 57%) and some non-coding genes (536, 7%) are represented by at least one known RefSeq transcript, labeled by the method “BestRefSeq” and assigned a transcript accession starting with NM_ or NR_, and corresponding RefSeq proteins designated with NP_ accessions. These are predominantly based on zebrafish mRNAs subject to manual and automated curation by the RefSeq team for over 20 years, including automated quality analyses and comparisons to previous zebrafish assemblies including GRCz11 (GCF_000002035.6) and ASM3317019v1 (GCA_033170195.1) to refine the annotations. 44% of the NM_ RefSeq transcripts included in RS_2025_04 have been fully reviewed by RefSeq curators.

Additional gene, transcript, and protein models were predicted using NCBI’s Gnomon algorithm using alignments of transcripts, proteins, and RNA-seq data as evidence. The evidence datasets used for RS_2025_04 are described at https://www.ncbi.nlm.nih.gov/refseq/annotation_euk/Danio_rerio/GCF_049306965.1-RS_2025_04/, and included alignments of available zebrafish mRNAs and ESTs, 6.4 billion RNA-seq reads from 710 SRA runs from a wide range of samples, 919 million 5’-Cap captured reads (CAGE or similar methods) from 66 SRA runs, 592 million PacBio or Oxford Nanopore transcript reads from 34 SRA runs, known RefSeq and GenBank proteins from zebrafish and other ray-finned fishes, and known RefSeq proteins from human. Additional non-coding models were generated using tRNAscanSE (v2.0.12), Infernal (v1.1.5) with Rfam v14.10 models for rRNAs and short ncRNAs, and miRNAs originally imported from miRBase release 22. Models from BestRefSeq, Gnomon, tRNAscan, and Infernal were combined to generate the final annotation, compared to the previous GCF_000002035.6-RS_2024_08 annotation of GRCz11 to retain GeneID, transcript, and protein accessions for equivalent annotations, and compared to the RefSeq annotation of human GRCh38 to identify orthologous genes. Gene nomenclature was based primarily on data from ZFIN^2^ and SwissProt proteins.

## References

1. Bradford, Y.M., Van Slyke, C.E., Howe, D.G., Fashena, D., Frazer, K., Martin, R., Paddock, H., Pich, C., Ramachandran, S., Ruzicka, L., et al. (2023). From multiallele fish to nonstandard environments, how ZFIN assigns phenotypes, human disease models, and gene expression annotations to genes. Genetics 224. 10.1093/genetics/iyad032.

2. Bradford, Y.M., Van Slyke, C.E., Ruzicka, L., Singer, A., Eagle, A., Fashena, D., Howe, D.G., Frazer, K., Martin, R., Paddock, H., et al. (2022). Zebrafish information network, the knowledgebase for research. Genetics 220. 10.1093/genetics/iyac016.

3. Varshney, G.K., Sood, R., and Burgess, S.M. (2015). Understanding and Editing the Zebrafish Genome. Adv Genet 92, 1–52. 10.1016/bs.adgen.2015.09.002.

4. Howe, K., Clark, M.D., Torroja, C.F., Torrance, J., Berthelot, C., Muffato, M., Collins, J.E., Humphray, S., McLaren, K., Matthews, L., et al. (2013). The zebrafish reference genome sequence and its relationship to the human genome. Nature 496, 498–503. 10.1038/nature12111.

5. Amores, A., Force, A., Yan, Y.L., Joly, L., Amemiya, C., Fritz, A., Ho, R.K., Langeland, J., Prince, V., Wang, Y.L., et al. (1998). Zebrafish hox clusters and vertebrate genome evolution. Science 282, 1711–1714. 10.1126/science.282.5394.1711.

6. Postlethwait, J., Amores, A., Force, A., and Yan, Y.L. (1999). The zebrafish genome. Methods Cell Biol 60, 149–163.

7. Collins, J.E., White, S., Searle, S.M.J., and Stemple, D.L. (2012). Incorporating RNA-seq data into the zebrafish Ensembl genebuild. Genome Res 22, 2067–2078. 10.1101/gr.137901.112.

8. Chernyavskaya, Y., Zhang, X.F., Liu, J.Z., and Blackburn, J. (2022). Long-read sequencing of the zebrafish genome reorganizes genomic architecture. Bmc Genomics 23. ARTN 116 10.1186/s12864-022-08349-3.

9. Song, J.Q., Dong, F.G., Lilly, J.W., Stupar, R.M., and Jiang, J.M. (2001). Instability of bacterial artificial chromosome (BAC) clones containing tandemly repeated DNA sequences. Genome 44, 463–469. DOI 10.1139/gen-44-3-463.

10. Eid, J., Fehr, A., Gray, J., Luong, K., Lyle, J., Otto, G., Peluso, P., Rank, D., Baybayan, P., Bettman, B., et al. (2009). Real-Time DNA Sequencing from Single Polymerase Molecules. Science 323, 133–138. 10.1126/science.1162986.

11. Jain, M., Koren, S., Miga, K.H., Quick, J., Rand, A.C., Sasani, T.A., Tyson, J.R., Beggs, A.D., Dilthey, A.T., Fiddes, I.T., et al. (2018). Nanopore sequencing and assembly of a human genome with ultra-long reads. Nat Biotechnol 36, 338-+. 10.1038/nbt.4060.

12. Nurk, S., Koren, S., Rhie, A., Rautiainen, M., Bzikadze, A.V., Mikheenko, A., Vollger, M.R., Altemose, N., Uralsky, L., Gershman, A., et al. (2022). The complete sequence of a human genome. Science 376, 44–53. 10.1126/science.abj6987.

13. Liu, J.L., Li, Q.L., Hu, Y.X., Yu, Y., Zheng, K., Li, D.F., Qin, L.X., and Yu, X.C. (2024). The complete telomere-to-telomere sequence of a mouse genome. Science 386, 1141–1146. 10.1126/science.adq8191.

14. Chen, J., Wang, Z.J., Tan, K.W., Huang, W., Shi, J.P., Li, T., Hu, J., Wang, K., Wang, C., Xin, B.B., et al. (2023). A complete telomere-to-telomere assembly of the maize genome. Nat Genet 55, 1221-+. 10.1038/s41588-023-01419-6.

15. Fodor, E., Okendo, J., Szabó, N., Szabó, K., Czimer, D., Tarján-Rácz, A., Szeverényi, I., Low, B.W., Liew, J.H., Koren, S., et al. (2024). The reference genome of Macropodus opercularis (the paradise fish). Sci Data 11. ARTN 540 10.1038/s41597-024-03277-1.

16. Sadamitsu, K., Velilla, F., Shinya, M., Kashima, M., Imai, Y., Kawasaki, T., Watai, K., Hosaka, M., Hirata, H., and Sakai, N. (2024). Establishment of a zebrafish inbred strain, M-AB, capable of regular breeding and genetic manipulation. Sci Rep-Uk 14. ARTN 7455 10.1038/s41598-024-57699-3.

17. Haffter, P., Granato, M., Brand, M., Mullins, M.C., Hammerschmidt, M., Kane, D.A., Odenthal, J., vanEeden, F.J.M., Jiang, Y.J., Heisenberg, C.P., et al. (1996). The identification of genes with unique and essential functions in the development of the zebrafish, Danio rerio. Development 123, 1–36.

18. Walker, C. (1999). Haploid screens and gamma-ray mutagenesis. Methods Cell Biol 60, 43–70. 10.1016/s0091-679x(08)61893-2.

19. Streisinger, G., Walker, C., Dower, N., Knauber, D., and Singer, F. (1981). Production of Clones of Homozygous Diploid Zebra Fish (Brachydanio-Rerio). Nature 291, 293–296. DOI 10.1038/291293a0.

20. Rautiainen, M., Nurk, S., Walenz, B.P., Logsdon, G.A., Porubsky, D., Rhie, A., Eichler, E.E., Phillippy, A.M., and Koren, S. (2023). Telomere-to-telomere assembly of diploid chromosomes with Verkko. Nat Biotechnol. 10.1038/s41587-023-01662-6.

21. Koren, S., Bao, Z.G., Guarracino, A., Ou, S.J., Goodwin, S., Jenike, K.M., Lucas, J., Mcnulty, B., Park, J., Rautiainen, M., et al. (2024). Gapless assembly of complete human and plant chromosomes using only nanopore sequencing. Genome Res 34, 1919–1930. 10.1101/gr.279334.124.

22. Rhie, A., Walenz, B.P., Koren, S., and Phillippy, A.M. (2020). Merqury: reference-free quality, completeness, and phasing assessment for genome assemblies. Genome Biol 21. ARTN 245 10.1186/s13059-020-02134-9.

23. Deng, Y., Qian, Y.T., Meng, M.H., Jiang, H.F., Dong, Y., Fang, C.C., He, S.P., and Yang, L.D. (2022). Extensive sequence divergence between the reference genomes of two zebrafish strains, Tuebingen and AB. Mol Ecol Resour 22, 2148–2157. 10.1111/1755-0998.13602.

24. Howe, K., Howard, C., McCarthy, S.A., Wood, J.M.D., Wellcome Sanger Institute Tree of Life Management, S., Laboratory, t., Wellcome Sanger Institute Scientific Operations: Sequencing, O., and Wellcome Sanger Institute Tree of Life Core Informatics, t. (2025). The genome sequence of the zebra danio, Danio rerio (Hamilton, 1822) (Cypriniformes: Danionidae). Wellcome Open Res 10, 330. 10.12688/wellcomeopenres.24569.1.

25. Showpnil, I.A., Gonzalez, M.E.H., Ramadesikan, S., Marhabaie, M., Daley, A., Dublin-Ryan, L., Pastore, M.T., Gurusamy, U., Hunter, J.M., Stone, B.S., et al. (2024). Long-read genome sequencing resolves complex genomic rearrangements in rare genetic syndromes. Npj Genom Med 9. ARTN 66 10.1038/s41525-024-00454-4.

26. Brown, K.H., Dobrinski, K.P., Lee, A.S., Gokcumen, O., Mills, R.E., Shi, X.H., Chong, W.W.S., Chen, J.Y.H., Yoo, P., David, S., et al. (2012). Extensive genetic diversity and substructuring among zebrafish strains revealed through copy number variant analysis. P Natl Acad Sci USA 109, 529–534. 10.1073/pnas.1112163109.

27. James, S.A., O’Kelly, M.J.T., Carter, D.M., Davey, R.P., van Oudenaarden, A., and Roberts, I.N. (2009). Repetitive sequence variation and dynamics in the ribosomal DNA array of as revealed by whole-genome resequencing. Genome Res 19, 626–635. 10.1101/gr.084517.108.

28. Nishibuchi, G., and Déjardin, J. (2017). The molecular basis of the organization of repetitive DNA-containing constitutive heterochromatin in mammals. Chromosome Res 25, 77–87. 10.1007/s10577-016-9547-3.

29. Vollger, M.R., Kerpedjiev, P., Phillippy, A.M., and Eichler, E.E. (2022). StainedGlass: interactive visualization of massive tandem repeat structures with identity heatmaps. Bioinformatics 38, 2049–2051. 10.1093/bioinformatics/btac018.

30. Li, K., Smith, M.L., Blazier, J.C., Kochan, K.J., Wood, J.M.D., Howe, K., Kwitek, A.E., Dwinell, M.R., Chen, H., Ciosek, J.L., et al. (2024). Construction and evaluation of a new rat reference genome assembly, GRCr8, from long reads and long-range scaffolding. Genome Res 34, 2081–2093. 10.1101/gr.279292.124.

31. Amores, A., and Postlethwait, J.H. (1999). Banded chromosomes and the zebrafish karyotype. Methods Cell Biol 60, 323–338. 10.1016/s0091-679x(08)61908-1.

32. Daga, R.R., Thode, G., and Amores, A. (1996). Chromosome complement, C-banding, Ag-NOR and replication banding in the zebrafish Danio rerio. Chromosome Res 4, 29–32. 10.1007/BF02254941.

33. Gornung, E., Gabrielli, I., Cataudella, S., and Sola, L. (1997). CMA3-banding pattern and fluorescence in situ hybridization with 18S rRNA genes in zebrafish chromosomes. Chromosome Res 5, 40–46. 10.1023/a:1018441402370.

34. Pijnacker, L.P., and Ferwerda, M.A. (1995). Zebrafish chromosome banding. Genome 38, 1052–1055. 10.1139/g95-140.

35. Sola, L., and Gornung, E. (2001). Classical and molecular cytogenetics of the zebrafish, Danio rerio (Cyprinidae, Cypriniformes): an overview. Genetica 111, 397–412. 10.1023/a:1013776323077.

36. de Sotero-Caio, C.G., Cabral-de-Mello, D., Calixto, M.D., Valente, G., Martins, C., Loreto, V., de Souza, M.J., and Santos, N. (2017). Centromeric enrichment of LINE-1 retrotransposons and its significance for the chromosome evolution of Phyllostomid bats. Chromosome Res 25, 313–325. 10.1007/s10577-017-9565-9.

37. Mashkova, T.D., Oparina, N.Y., Lacroix, M.H., Fedorova, L.I., Tumeneva, I.G., Zinovieva, O.L., and Kisselev, L.L. (2001). Structural rearrangements and insertions of dispersed elements in pericentromeric alpha satellites occur preferably at kinkable DNA sites. J Mol Biol 305, 33–48. 10.1006/jmbi.2000.4270.

38. Logsdon, G.A., Rozanski, A.N., Ryabov, F., Potapova, T., Shepelev, V.A., Catacchio, C.R., Porubsky, D., Mao, Y.F., Yoo, D., Rautiainen, M., et al. (2024). The variation and evolution of complete human centromeres. Nature 629. 10.1038/s41586-024-07278-3.

39. Chen, J.L., Blasco, M.A., and Greider, C.W. (2000). Secondary structure of vertebrate telomerase RNA. Cell 100, 503–514. 10.1016/s0092-8674(00)80687-x.

40. Lin, Y.L., and Gokcumen, O. (2019). Fine-Scale Characterization of Genomic Structural Variation in the Human Genome Reveals Adaptive and Biomedically Relevant Hotspots. Genome Biol Evol 11, 1136–1151. 10.1093/gbe/evz058.

41. Goldfarb, T., Kodali, V.K., Pujar, S., Brover, V., Robbertse, B., Farrell, C.M., Oh, D.H., Astashyn, A., Ermolaeva, O., Haddad, D., et al. (2024). NCBI RefSeq: reference sequence standards through 25 years of curation and annotation. Nucleic Acids Res 53, D243–D257. 10.1093/nar/gkae1038.

42. Monnich, M., Borgeskov, L., Breslin, L., Jakobsen, L., Rogowski, M., Doganli, C., Schroder, J.M., Mogensen, J.B., Blinkenkjaer, L., Harder, L.M., et al. (2018). CEP128 Localizes to the Subdistal Appendages of the Mother Centriole and Regulates TGF-beta/BMP Signaling at the Primary Cilium. Cell Rep 22, 2584–2592. 10.1016/j.celrep.2018.02.043.

43. Vollger, M.R., Guitart, X., Dishuck, P.C., Mercuri, L., Harvey, W.T., Gershman, A., Diekhans, M., Sulovari, A., Munson, K.M., Lewis, A.P., et al. (2022). Segmental duplications and their variation in a complete human genome. Science 376, 55-+. ARTN eabj6965 10.1126/science.abj6965.

44. Samonte, R.V., and Eichler, E.E. (2002). Segmental duplications and the evolution of the primate genome. Nat Rev Genet 3, 65–72. DOI 10.1038/nrg705.

45. Emanuel, B.S., and Shaikh, T.H. (2001). Segmental duplications: An ’expanding’ role in genomic instability and disease. Nat Rev Genet 2, 791–800. Doi 10.1038/35093500.

46. Hicktey, G., Monslong, J., Ebler, J., Novak, A.M., Eizenga, J.M., Gao, Y., Marschall, T., Li, H., Paten, B., Abel, H.J., et al. (2024). Pangenome graph construction from genome alignments with Minigraph-Cactus. Nat Biotechnol 42. 10.1038/s41587-023-01793-w.

47. Guarracino, A., Heumos, S., Nahnsen, S., Prins, P., and Garrison, E. (2022). ODGI: understanding pangenome graphs. Bioinformatics 38, 3319–3326. 10.1093/bioinformatics/btac308.

48. Suurväli, J., Whiteley, A.R., Zheng, Y.C., Gharbi, K., Leptin, M., and Wiehe, T. (2020). The Laboratory Domestication of Zebrafish: From Diverse Populations to Inbred Substrains. Mol Biol Evol 37, 1056–1069. 10.1093/molbev/msz289.

49. Honjo, Y., Takano, K., and Ichinohe, T. (2021). Characterization of novel zebrafish MHC class I U lineage genes and their haplotype. Dev Comp Immunol 116. ARTN 103952 10.1016/j.dci.2020.103952.

50. Ebert, P., Audano, P.A., Zhu, Q., Rodriguez-Martin, B., Porubsky, D., Bonder, M.J., Sulovari, A., Ebler, J., Zhou, W., Serra Mari, R., et al. (2021). Haplotype-resolved diverse human genomes and integrated analysis of structural variation. Science 372. 10.1126/science.abf7117.

51. Sherman, R.M., Forman, J., Antonescu, V., Puiu, D., Daya, M., Rafaels, N., Boorgula, M.P., Chavan, S., Vergara, C., Ortega, V.E., et al. (2018). Assembly of a pan-genome from deep sequencing of 910 humans of African descent. Nat Genet 51, 30–35. 10.1038/s41588-018-0273-y.

52. Baranasic, D., Hortenhuber, M., Balwierz, P.J., Zehnder, T., Mukarram, A.K., Nepal, C., Varnai, C., Hadzhiev, Y., Jimenez-Gonzalez, A., Li, N., et al. (2022). Multiomic atlas with functional stratification and developmental dynamics of zebrafish cis-regulatory elements. Nat Genet 54, 1037–1050. 10.1038/s41588-022-01089-w.

53. Varga, Z.M. (2016). Aquaculture, husbandry, and shipping at the Zebrafish International Resource Center. Methods Cell Biol 135, 509–534. 10.1016/bs.mcb.2016.01.007.

54. Varga, Z.M., and Murray, K.N. (2016). Health monitoring and disease prevention at the Zebrafish International Resource Center. Methods Cell Biol 135, 535–551. 10.1016/bs.mcb.2016.04.020.

55. Murray, K.N., Varga, Z.M., and Kent, M.L. (2016). Biosecurity and Health Monitoring at the Zebrafish International Resource Center. Zebrafish 13 *Suppl 1*, S30–38. 10.1089/zeb.2015.1206.

56. Matthews, J.L., Murphy, J.M., Carmichael, C., Yang, H., Tiersch, T., Westerfield, M., and Varga, Z.M. (2018). Changes to Extender, Cryoprotective Medium, and In Vitro Fertilization Improve Zebrafish Sperm Cryopreservation. Zebrafish 15, 279–290. 10.1089/zeb.2017.1521.

57. Westerfield, M. (2007). The Zebrafish Book. A Guide for the Laboratory Use of Zebrafish (Danio rerio), 5th Edition Edition (University of Oregon Press).

58. Rautiainen, M., and Marschall, T. (2020). GraphAligner: rapid and versatile sequence-to-graph alignment. Genome Biol 21. ARTN 253 10.1186/s13059-020-02157-2.

59. Goel, M., Sun, H.Q., Jiao, W.B., and Schneeberger, K. (2019). SyRI: finding genomic rearrangements and local sequence differences from whole-genome assemblies. Genome Biol 20. ARTN 277 10.1186/s13059-019-1911-0.

60. Porubsky, D. (2024). SVbyEye: A visual tool to characterize structural variation among whole-genome assemblies. 10.1101/2024.09.11.612418.

