## supplementary figures for "Complete *de novo* assembly and re-annotation of the zebrafish genome"

#### Summary of Figures

**Supplementary Fig. 9:** Structural variation comparisons between assemblies. SVbyEye [19] plot showing the structural variations following the comparative analysis of the GRCz12tu and the GRCz12ab genome assemblies. The “Target sequence” is GRCz12tu and the query sequence is the GRCz12ab assembly. The red on the plot denotes deletions the blue highlights the insertion locations; green color represent the forward orientation of the chromosomes between the assemblies. The centromere is marked by black. ........ 17

**Supplementary Fig. 10:** Chromosomal comparisons of zebrafish centromere lengths from the GRCz12ab (red), GRCz12tu (green), and NHGRI_Fish11 (blue) assemblies. Each point on the plot represents the centromere lengths of a given chromosome on the x-axis. The lower plot shows the zebrafish strain differences in centromere lengths. .... 18

**Supplementary Fig. 11:** GRCz11 and GRCz12tu ribosomal RNA gene copy number quantifications. The bar plot shows the gene copy numbers from the two assemblies we compared. Chromosome 4 (NC_007115.7) is highly enriched with all rRNA genes compared with the rest of the chromosomes in GRCz12tu and GRCz11 assemblies. .. 19

**Supplementary Fig. 12:** Distribution of repetitive elements across the centromeres of selected chromosomes of the zebrafish and Macropodus opercularis (paradise fish). The plot displays the centromeric distribution of non-satellite insertions in GRCz12tu and macOpe2 genomes. Each row represents the position and the orientation of the repeat element along the centromeres, and the color coding denotes the repeats of identity. Chromosome numbers in MacOpe2 do not correlate to the same chromosomes in GRCz12tu........................................................................................................ 20

**Supplementary Fig. 13: Linear visualization of pangenome graph using ODGI[22].** The pangenome graph was generated with odgi viz, showing the alignment and structural variation across 24 zebrafish. Each horizontal-colored bar graph represents a haplotype-specific path traversing the graph. Vertical disruptions represent variations such as deletions and insertions. The black lines show the underlying graph topology. The consistent horizontal patterns reflect conserved regions, while the “breaks” shows regions of complexity in the zebrafish genomes. Pangenome graph showing the structural and gene copy number variation in zebrafish chromosomes 19 and 24, respectively. ......... 21

**Supplementary Fig. 14:** Principal Component Analysis (PCA) plot displaying the genetic variation among the zebrafish we compared. The plot was rendered using variants obtained from the pangenome multi-sample VCF file and it reveals substantial genetic differences where each point represents the individual fish. The wild-caught EKK and CB fish cluster most strongly with each other while AB and TU are more related to each other despite being obtained from presumably independent sources. The GRCz12tu is missing in the plot because it was used as a reference and it does not get included in the final VCF we used in building PCA. .......................................................................... 22

### Materials and methods

#### Library preparation and sequencing

##### Oxford Nanopore Technology (ONT), ultra-long sequencing

GenomicDNA libraries were created from ultra-high molecular weight DNA from zebrafish cell culture using the Oxford Nanopore sequencing kit SQK-ULK114. Each library was loaded onto a FLO-PRO114M flow cell and run for 72 hours or more with two subsequent loadings at 24-hour intervals. The libraries were sequenced on a PromethION (Oxford Nanopore) running MinKNOW software version MinKNOW 24.02.19. Base calling and modified base detection (5mC, 5hmC) were performed using Dorado 7.3.11.

##### PacBio HiFi sequencing

The Pacific Biosciences protocol “Preparing whole genome and metagenome libraries using SMRTbell® prep kit 3.0” was used to create a library from 15 micrograms of DNA. The Megarupter (Diagenode) was used for shearing and SageELF (Sage Science) was used for size-selection for fragments ~20 kb. Library size was assessed using a FemtoPulse (Agilent). Each library was run on 25M SMRTCells using Revio sequencing reagents. Sequencing was performed on a Revio sequencer (Pacific Biosciences) running instrument control version 13.1.0.221972 and a movie collection time of 30 hours per SMRTCell with a 2-hour pre-extension and adaptive loading. CCS/HiFi unaligned bam files with kinetic information and 5mC tags (called with jasmine) were generated on-instrument using tools in software version 13.1.0.221972. Fastq sequences were extracted from the bam files using bam2fastq (version 3.2.0 / SMRTLink 13.1.0.221970).

##### PacBio Iso-Seq library preparation and sequencing

Iso-Seq libraries were created using the PacBio protocol “Preparing Kinnex™ libraries using the Kinnex full-length RNA kit” protocol (Pacific Biosciences) from 300 ng of RNA. Each library was run on version 25M using Revio sequencing reagents. Sequencing was performed on a Revio sequencer (Pacific Biosciences) running instrument control version 13.1.0.221972 with a movie collection time of 30 hours, 1.42-hour pre-extension, and adaptive loading. Circular consensus sequence (CCS) reads were generated by the Revio on-instrument software. Transcript arrays in CCS reads were deconcatenated with skera (version 1.2.0) to generate segmented reads (Sreads). To generate full-length (FL) CCS reads, Sreads with the proper orientation of 5' and 3' Iso-Seq primers on the sequence ends were identified using lima (version 2.10.0). Full-length non-chimeric (FLNC) reads were generated from the FL reads using isoseq (version 4.1.2) refine which identifies FL reads with the correct 5prime and 3prime barcode orientation and filters out concatemerized products.

### Assembly

#### Assembly orientation

During the assembly, RagTag version 2.1.0[1] was used to orient the T2T assemblies using GRCz11 as a reference. The following command was used to orient the T2T chromosomes: *ragtag.py scaffold GRCz11.fasta NHGRI_Fish6_T2T.fasta -o ragtagoutput -t 24 -w -C.*

#### Polishing and genome assembly validation

The resulting genome assemblies were polished by followed the procedures previously used by Rhie et al [2]. Specifically, errors resembling short nucleotide variations (SNVs) were identified using DeepVariant version 1.8.0[3] from PacBio HiFi long reads, then filtered with BCFtools[4] and Merfin[5] to correct consensus and phasing errors. The assemblies, both before and after polishing, were assessed using k-mers derived from PacBio HiFi reads with Merqury version 1.4[6].

#### Genome coverage analysis

The Oxford Nanopore Technology (ONT), and PacBio circular consensus sequencing (HiFi) reads were mapped back to the assembled genome to examine consensus coverage and accuracy. HiFi and ONT sequencing data were aligned to assembled genome using winnowmap2 version 2.0.3[7]. The T2T-polishing (https://github.com/arangrhie/T2T-Polish) procedure was used to generate the coverage statistics. The "*cov.bed*" file output from the T2T polish script was then used to plot the chromosomal genome coverage using R programming language.

#### Quality value (QV) calculation of the assembled genome

A 21-mer database was constructed from the PacBio HiFi sequencing data using Meryl which is part of the Merqury tool suite. Assembly quality value (QV) and the error rate assembly statistics for the final polished genome assembly was assed using Merqury version 1.4[6]. We achieved genomic QV of 60 and the chromosomal QVs ranged from 50.53 to 62.60 for GRCz12tu (supplementary Table S1). For NIHGRI_Fish11 genomic QV was 52 and the chromosomal QV ranged from 42.65 to 61.29 (supplementary Table S2). The GRCz12ab achieved a chromosomal QV value of 59.72 with chromosomal QV scores that ranged from 53.7 to 63.1 (Supplementary Table S3).

### Structural variation and syntenic analysis

The structural variation analysis was conducted by aligning the NHGRI_Fish11 to GRCz12tu and or the GRCz12ab assembly to GRCz12tu using minimap2 [8] and sorted with samtools[9] using the following options: *minimap2 -ax asm20 -t 24 --eqx GRCz12tu.fasta GRCz12ab.fasta | samtools sort -O BAM - > output.bam* . The structural variation was then done using Sniffles2[10] version 2.6.1 using the *output.bam* as the input. The resulting structural variant types in the Variants Call Format (VCF) file were then summarized in R. To visualize the syntenic blocks between our assemblies and GRCz11, the GRCz12tu, NHGRI_Fish11, and GRCz12ab genome assemblies were aligned to GRCz11 using minimap2[8] with the following options: *minimap2 -x asm5 -c --eqx --secondary=no -t 24 -o output.paf GRCz12tu.fasta NHGRI_Fish11.fasta.* The resulting Sequence Alignment Map (SAM) file was then converted to Binary Alignment Map (BAM) and then indexed using samtools[9]. Syntenic and Rearrangement Identifier (SyRi)[11] was used to detect synteny blocks between assemblies and GRCz11.Plotstr was then used to visualize the syntenic blocks and all presumed structural variations between GRCz11,GRCz12tu, GRCz12ab, and NHGRI_Fish11 assemblies.

##### Identifying previously unassembled regions

To identify the previously unassembled regions, the GRCz11 was aligned to the complete GRCz12tu genome assembly using winnowmap2 [7]. The sequences that did not map to the GRCz12tu genome were then extracted using the following method:

cat winnowmap.paf | awk '{if ($12 > 0) print $6\t$8\t$9}' | bedtools sort -I - | bedtools merge -I - | bedtools complement -I - -g chr.len > pur.regions

The pur.regions contained the coordinates of the previously unassembled regions of the genomes which were then extracted using "bedtools getfasta -fi GRCz12tu.fasta -bed pur.regions -fo pur.fasta"

#### Identification of segmental duplications

Tandem repeat sequences in the final, assembled genomes were annotated using Tandem Repeats Finder (TRF)[12] and RepeatMasker[13]. Segmental duplications (SDs) were identified with biser v1.4[14] on the soft-masked genomes, employing the following parameters:

*biser -t 30 --max-error 20 --max-edit-error 10 --output GRcz12tu.sd --gc-heap 20G --kmer-size 31 GRCz12tu.fasta. soft.masked.* To identify the functional SDs, we considered the SDs with length equal or greater than 20kbs and are at least 500kb apart.

##### Centromere repeats identification

Using the GRCz12tu and GRCz12ab genome assemblies, we used the CentroMiner, part of the quarTeT tool suite[15] to predict centromere-associated repetitive sequences, such as satellite repeats and higher-order repeats (HORs). Based on repeat density, sequence homogeneity, and self-similarity patterns, the tool identified candidate centromeric regions. We manually checked and selected regions with the highest enrichment of satellite repeats. We then used StainedGlass[16] to confirm CentroMiner's centromere enrichment coordinates. The centromere sequences were then extracted using "bedtools getfasta"[17] using centromere coordinates in BED format. The identification and characterization of non-satellites insertions in the centromere was then done using EDTA software[18] with the centromere fasta file from the "bedtools getfasta" command. SVbyEye[19] was then used to annotate the chromosomal location of centromeres on the selected chromosomes.

#### Pangenome construction and analysis

We assembled draft genomes from twenty different wild-caught zebrafish using Verkko [20]. We then built the pangenome graph using Minigraph-Cactus pipeline[21] using the twenty different wild-caught zebrafish (40 haplotypes) plus the 3 complete assemblies and fDanRer4.1 laboratory strains and the downstream analysis was done using ODGI [22]. The use of Minigraph-Cactus enabled the accurate representation of structural variations between the 44 haplotype assemblies that we compared.

### Supplementary figures

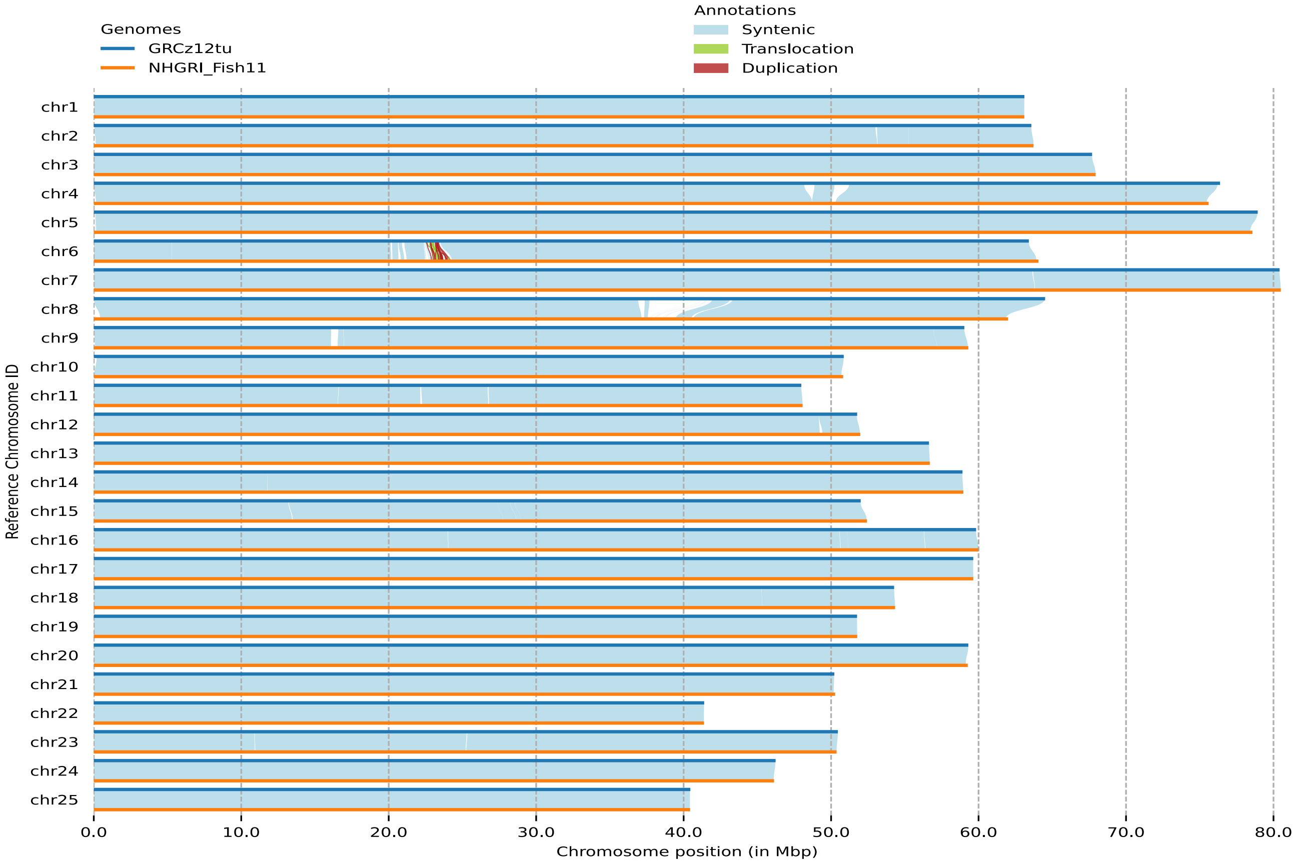

**Supplementary Fig 1:** SyRi syntenic plot of GRCz12tu and NHGRI_Fish11 genome assemblies. The large, segmental differences between the assemblies is highlighted as translocation and duplications.

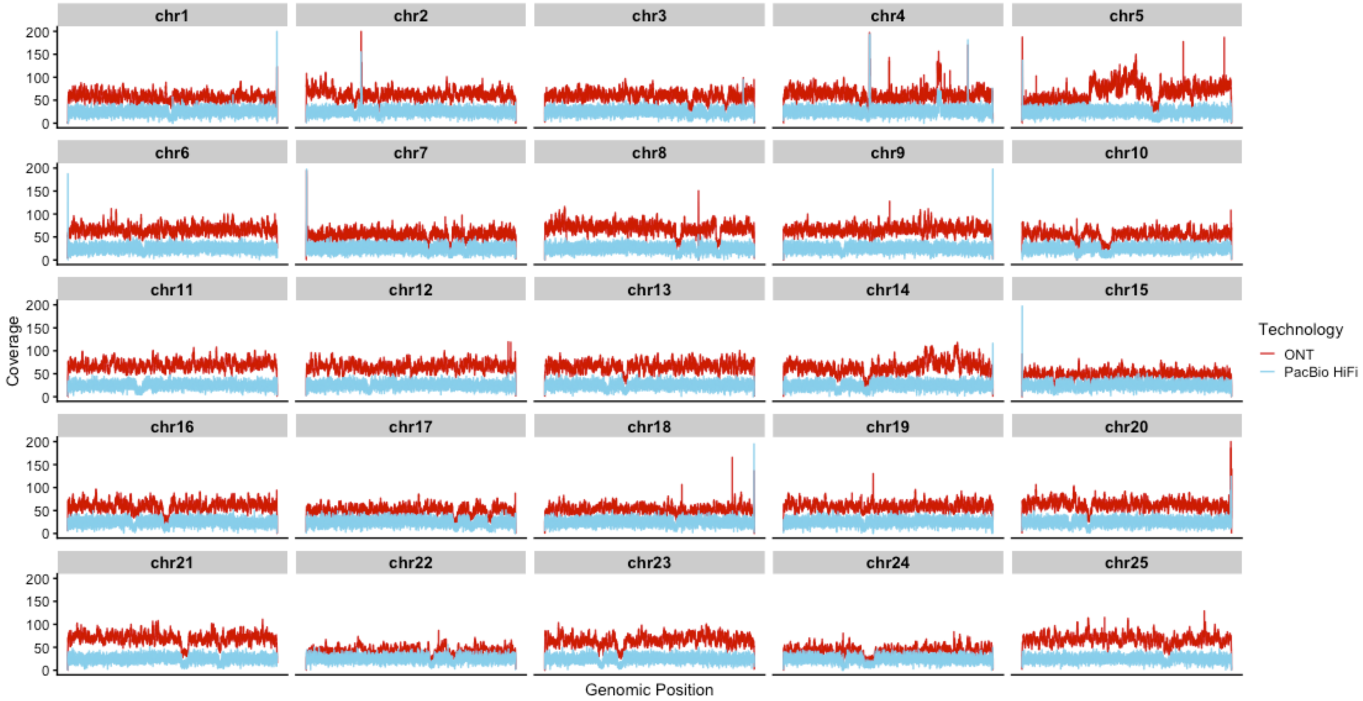

**Supplementary Fig 2:** GRCz12tu genome coverage plot showing the sequence depth coverage of ONT and PacBio HiFi reads mapping to the GRCz12tu genome assembly. ONT (red) and PacBio HiFi (blue) reads coverage across the 25 chromosomes in GRCz12tu genome assembly. The mapping exhibited uniform coverage across the genome and the dips in coverage corresponded to the centromeric location and rDNA arrays.

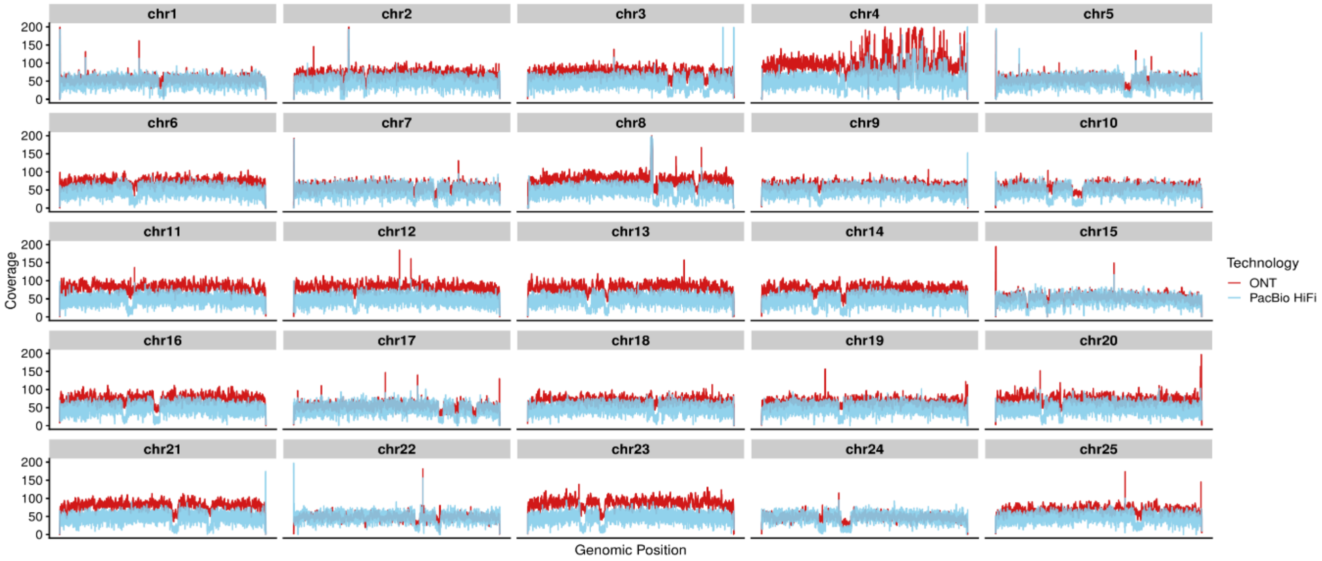

**Supplementary Fig 3:** Mapping coverage of the NHGRI_Fish11 genome assembly. Red and blue colors correspond to ONT and PacBio HiFi mapping coverages, respectively.

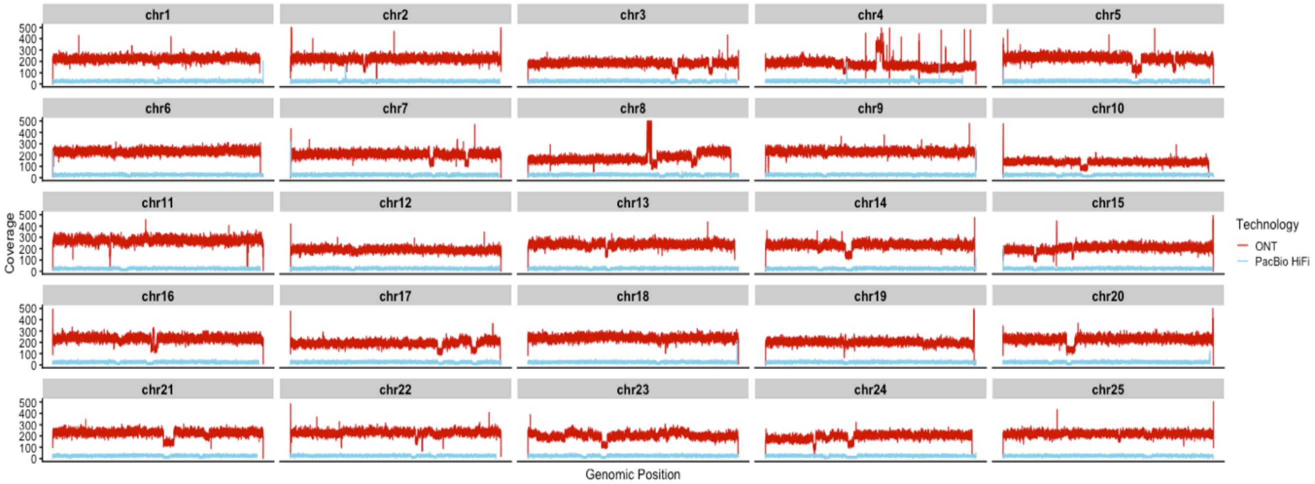

**Supplementary Fig 4:** GRCz12ab genome assembly ONT and PacBio genome coverage plot. The blue color in the figure legend represent the PacBio HiFi reads coverage, and the red color denotes the ONT mapping coverage.

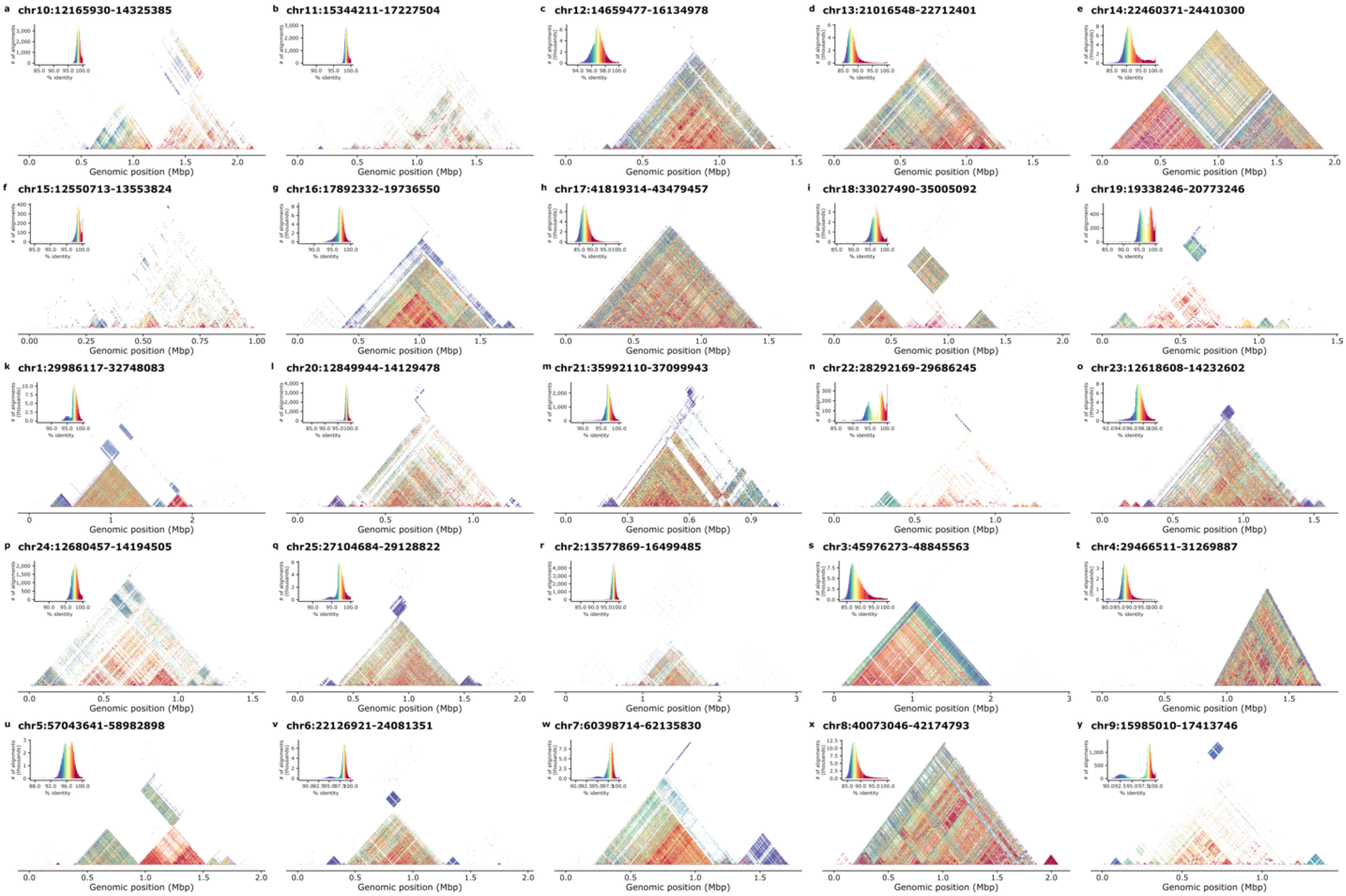

**Supplementary Fig 5:** Identification of chromosomal centromere repeats and locations. Centromere identity heatmaps produced using StainedGlass [16] display centromere locations across each chromosome. The regions with low identity suggest the presence of non-satellite sequence insertions.

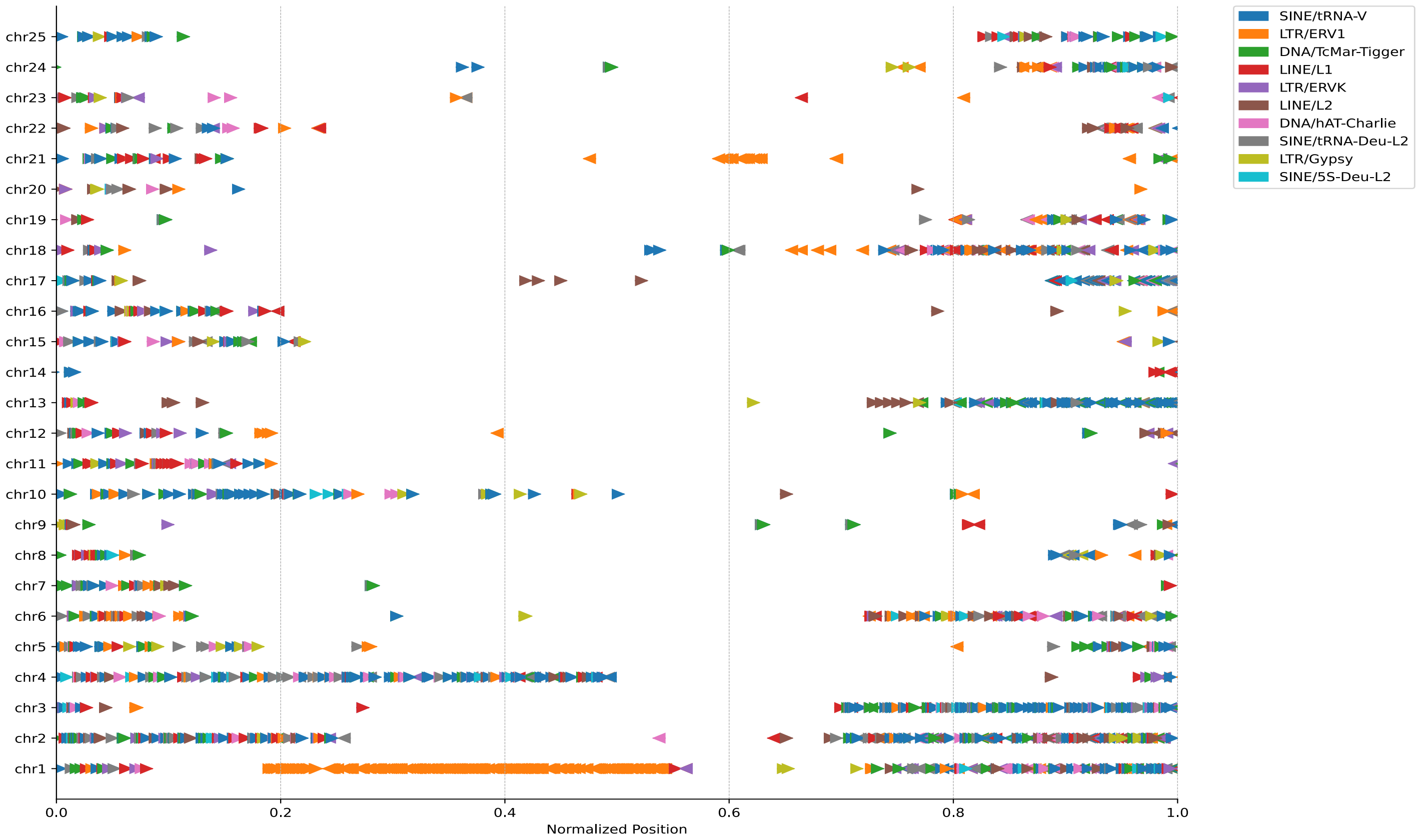

**Supplementary Fig 6:** Centromeric distribution and orientation of non-satellite insertions in centromeres of GRCz12tu. The identity of non-satellite insertions is shown in the top-right corner of the figure.

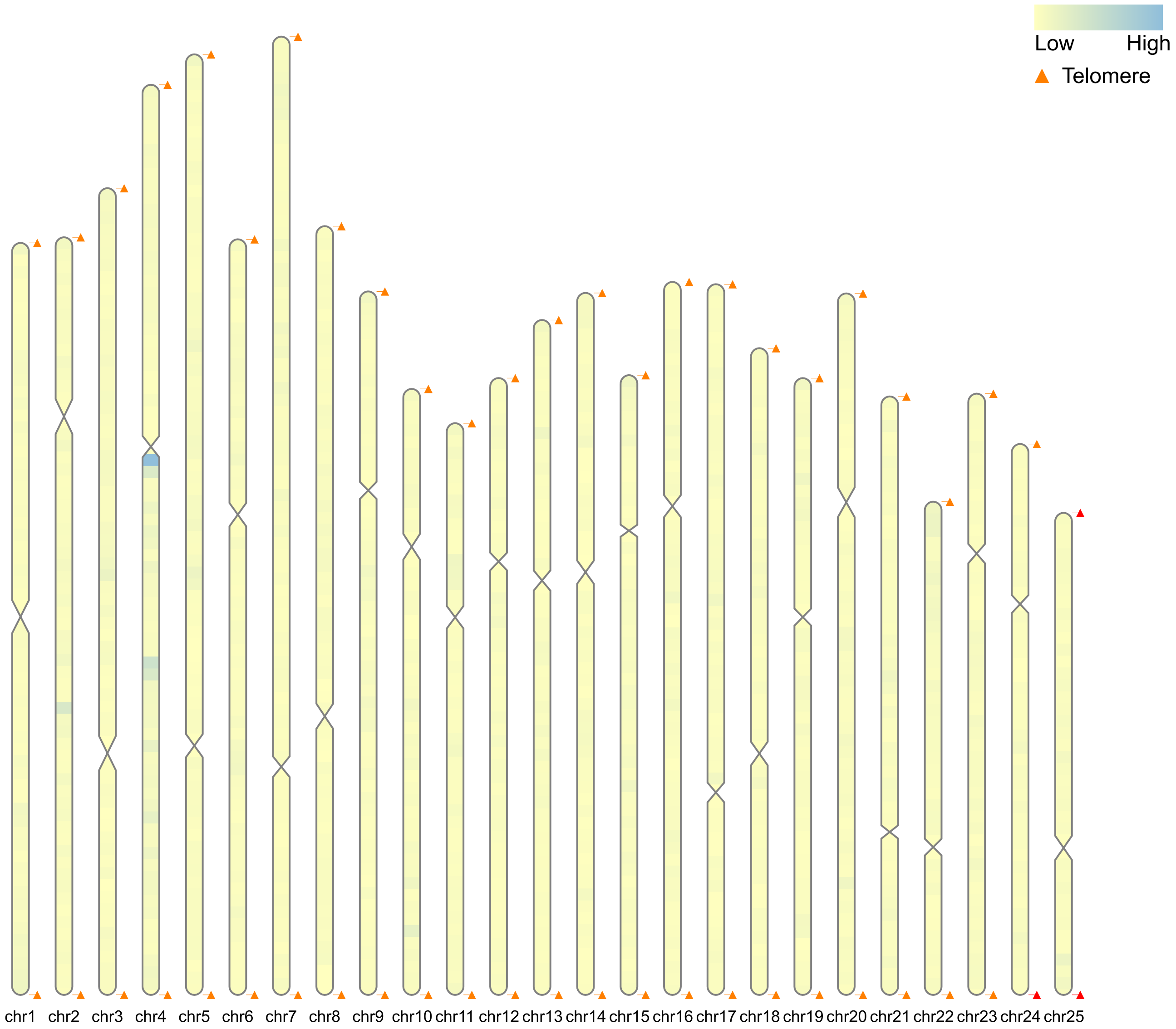

**Supplementary Fig 7:** Karyotype plot of the GRCz12tu genome showing all the chromosomal locations of the telomeric hexamer sequences. The constrictions are the centromere locations for all 25 chromosomes.

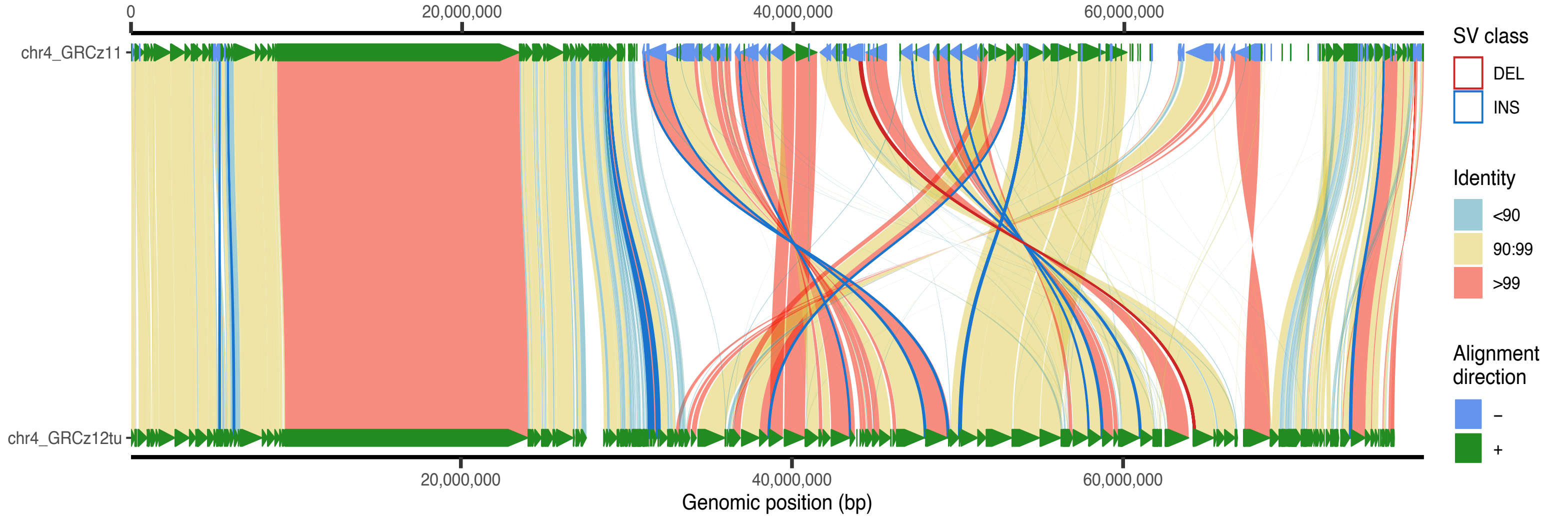

**Supplementary Fig 8:** Syntenic analysis showing the resolved chromosome 4. The alignment direction shows both forward (green) and reverse (blue) regions of the GRCz11 and GRCz12tu assemblies.

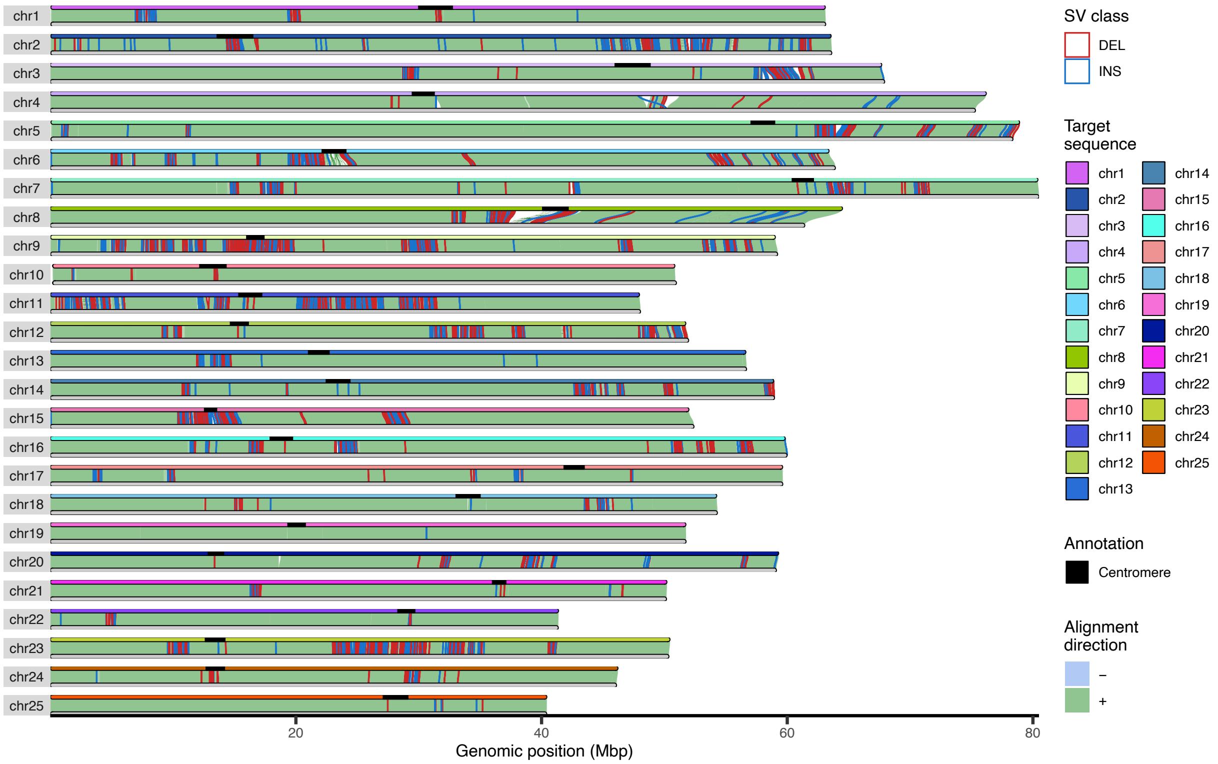

**Supplementary Fig 9:** Structural variation comparisons between assemblies. SVbyEye [19] plot showing the structural variations following the comparative analysis of the GRCz12tu and the GRCz12ab genome assemblies. The "Target sequence" is GRCz12tu and the query sequence is the GRCz12ab assembly. The red on the plot denotes deletions the blue highlights the insertion locations; green color represents the forward orientation of the chromosomes between the assemblies. The centromere is marked by black.

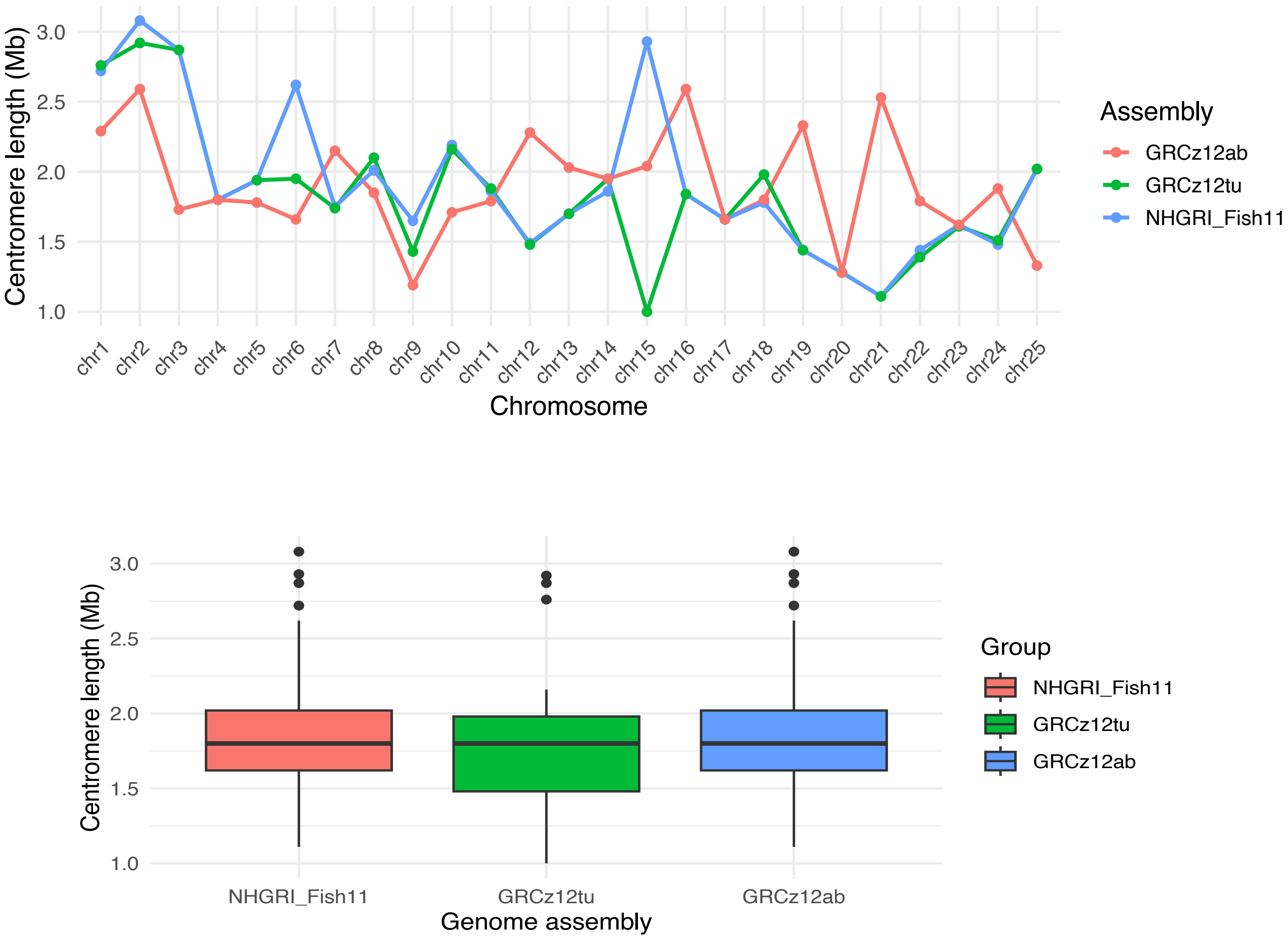

**Supplementary Fig. 10:** Chromosomal comparisons of zebrafish centromere lengths from the GRCz12ab (red), GRCz12tu (green), and NHGRI_Fish11 (blue) assemblies. Each point on the plot represents the centromere lengths of a given chromosome on the x-axis. The lower plot shows the zebrafish strain differences in centromere lengths.

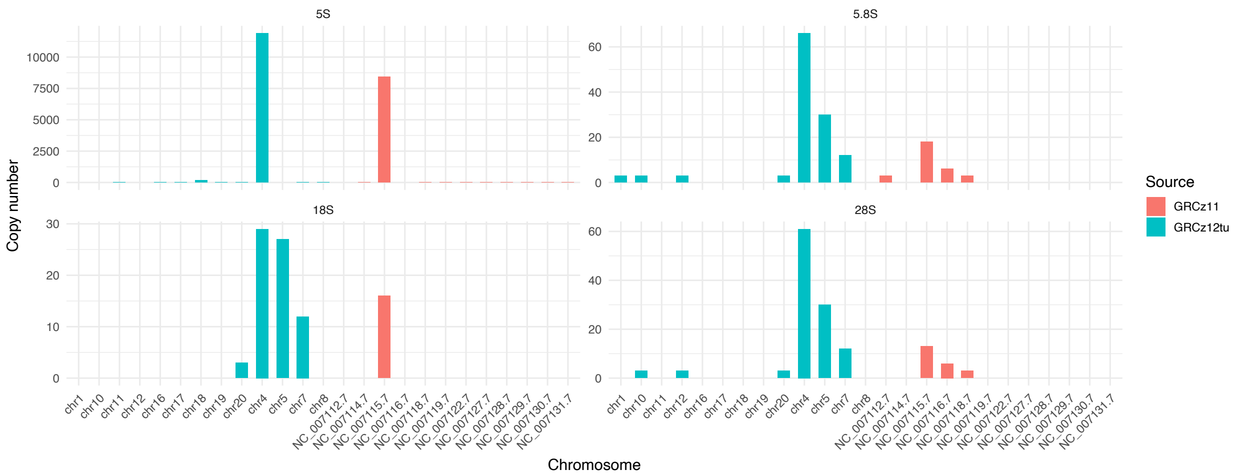

**Supplementary Fig. 11:** GRCz11 and GRCz12tu ribosomal RNA gene copy number quantifications. The bar plot shows the gene copy numbers from the two assemblies we compared. Chromosome 4 (NC_007115.7) is highly enriched with all rRNA genes compared with the rest of the chromosomes in GRCz12tu and GRCz11 assemblies.

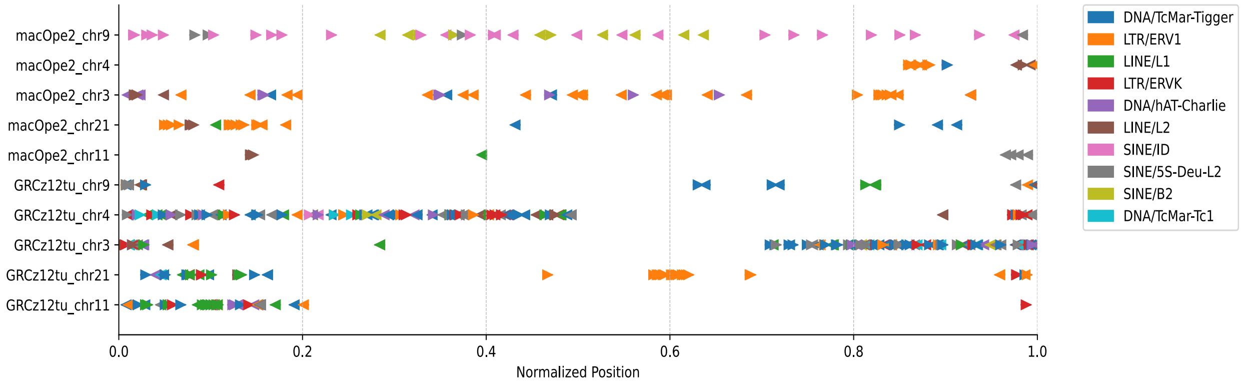

**Supplementary Fig. 12:** Distribution of repetitive elements across the centromeres of selected chromosomes of the zebrafish and *Macropodus opercularis* (paradise fish). The plot displays the centromeric distribution of non-satellite insertions in GRCz12tu and macOpe2 genomes. Each row represents the position and the orientation of the repeat element along the centromeres, and the color coding denotes the repeats of identity. Chromosome numbers in MacOpe2 do not correlate to the same chromosomes in GRCz12tu.

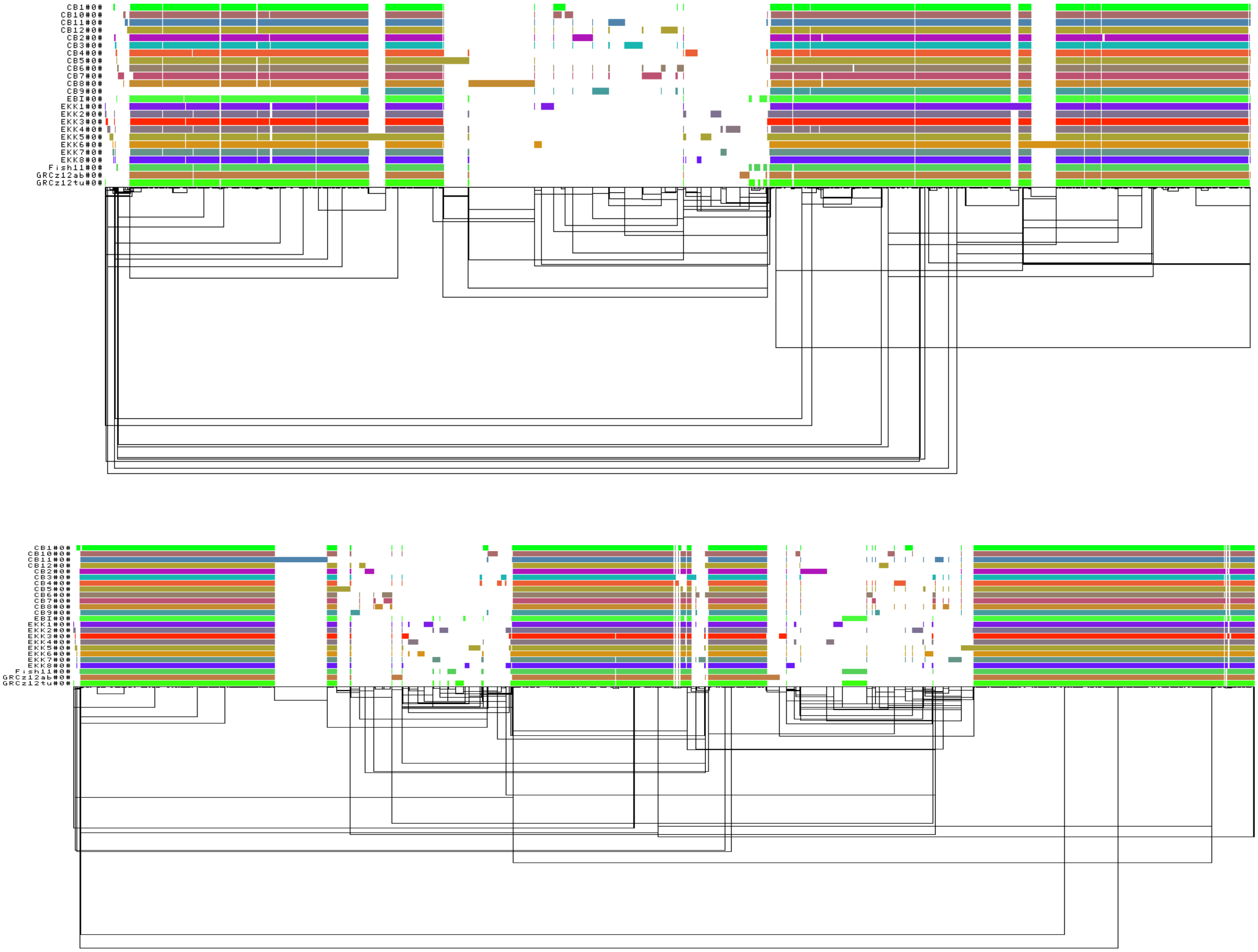

**Supplementary Fig 13: Linear visualization of pangenome graph using ODGI[22].** The pangenome graph was generated with odgi viz, showing the alignment and structural variation across 24 zebrafish. Each horizontal-colored bar graph represents a haplotype-specific path traversing the graph. Vertical disruptions represent variations such as deletions and insertions. The black lines show the underlying graph topology. The consistent horizontal patterns reflect conserved regions, while the "breaks" shows regions of complexity in the zebrafish genomes. Pangenome graph showing the structural and gene copy number variation in zebrafish chromosomes 19 and 24, respectively.

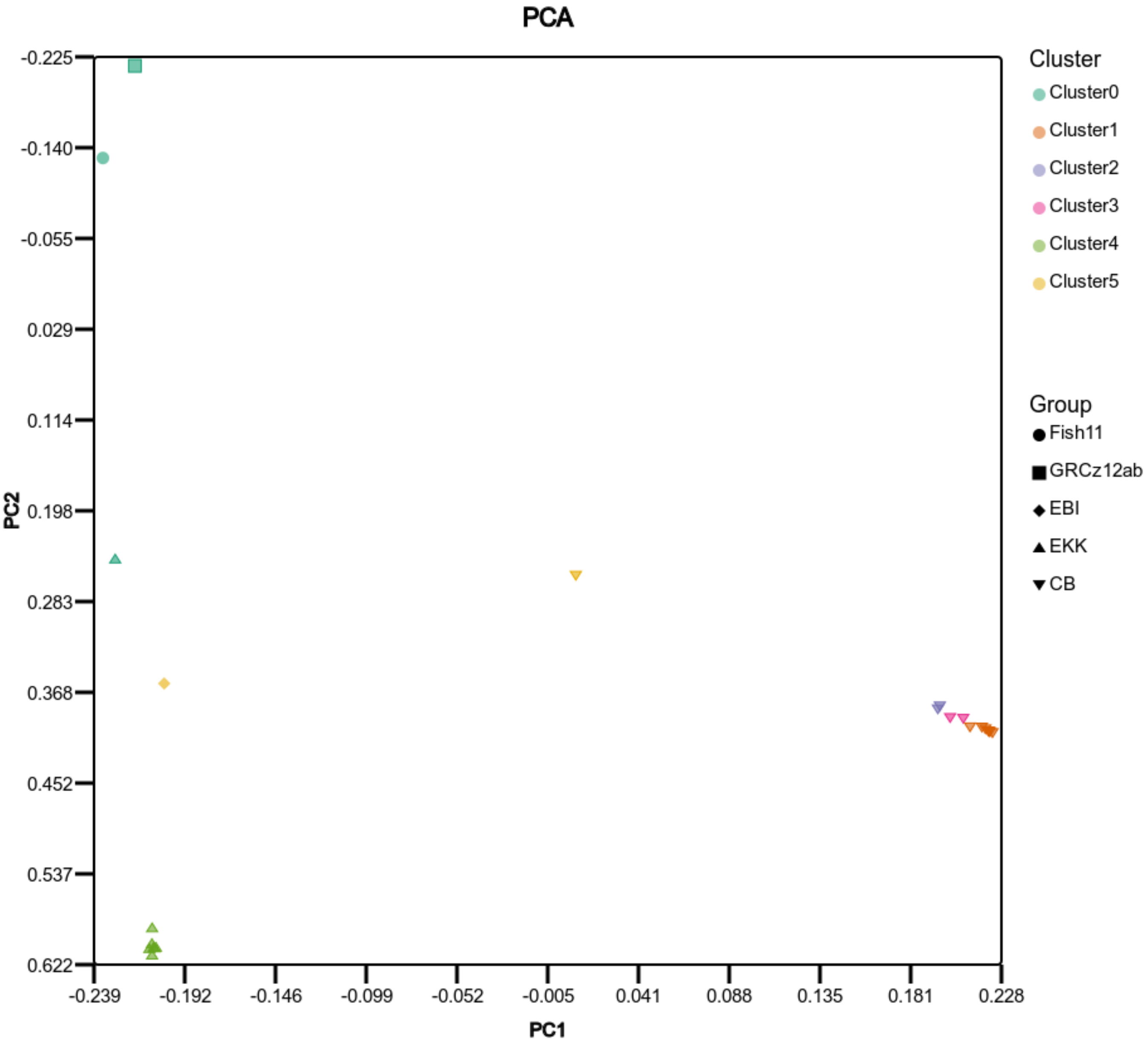

**Supplementary Fig14:** Principal Component Analysis (PCA) plot displaying the genetic variation among the zebrafish we compared. The plot was rendered using variants obtained from the pangenome multi-sample VCF file and it reveals substantial genetic differences where each point represents the individual fish. The wild-caught EKK and CB fish cluster most strongly with each other while AB and TU are more related to each other despite being obtained from presumably independent sources. The GRCz12tu was used as a reference therefore does not get included in the final VCF used in building the PCA.

**A**

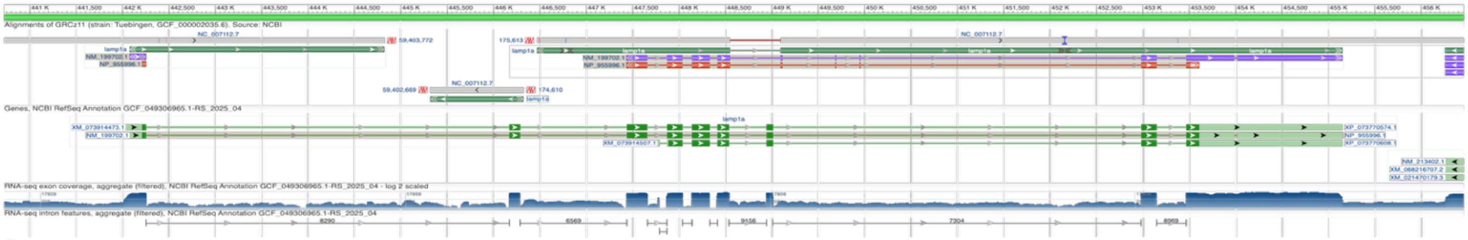

**B**

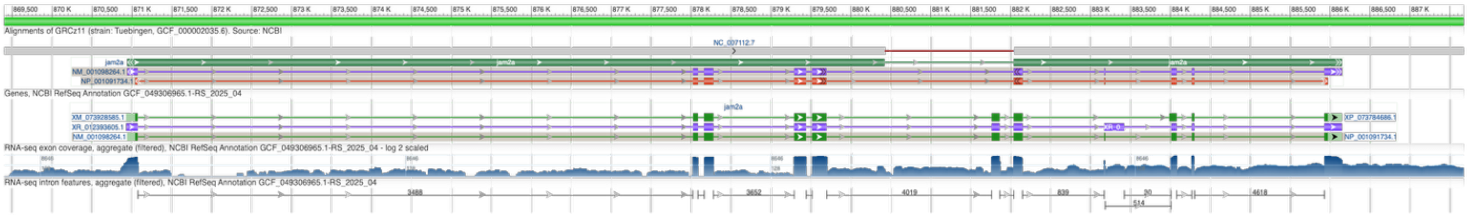

**C**

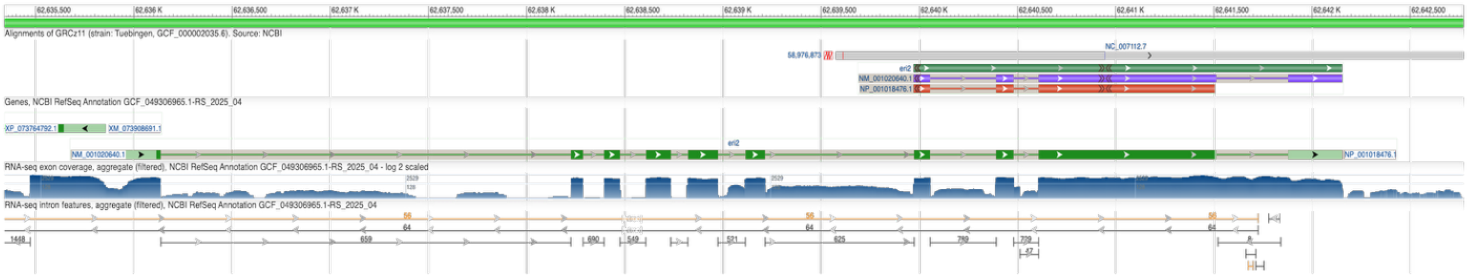

**Supplementary Fig 15:** Repair of incorrect gene models in GRCz12tu. (**A**) is the resolution of the missing coding sequences in the *lamp1* gene in GRCz11, (**B**) The *jam2a* gene was lacking coding sequence in GRCz11 which is now corrected (**C**), *eri2* was missing six coding sequences and is now fully resolved in GRCz12tu.

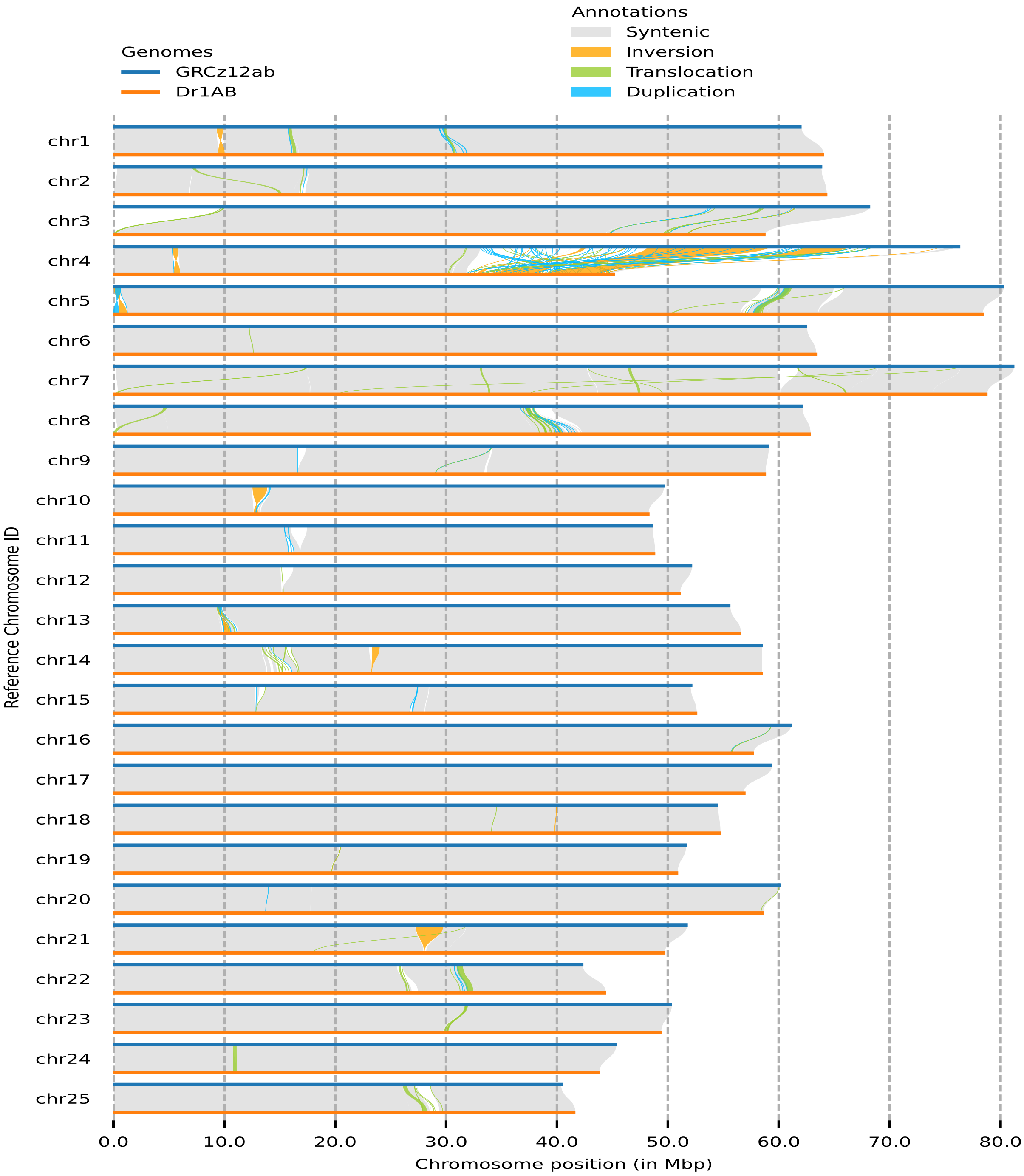

**Supplementary Fig 16:** Syntenic comparison of GRCz12ab and Dr1AB genome assemblies. The annotation shows the segmental differences between the two assemblies.

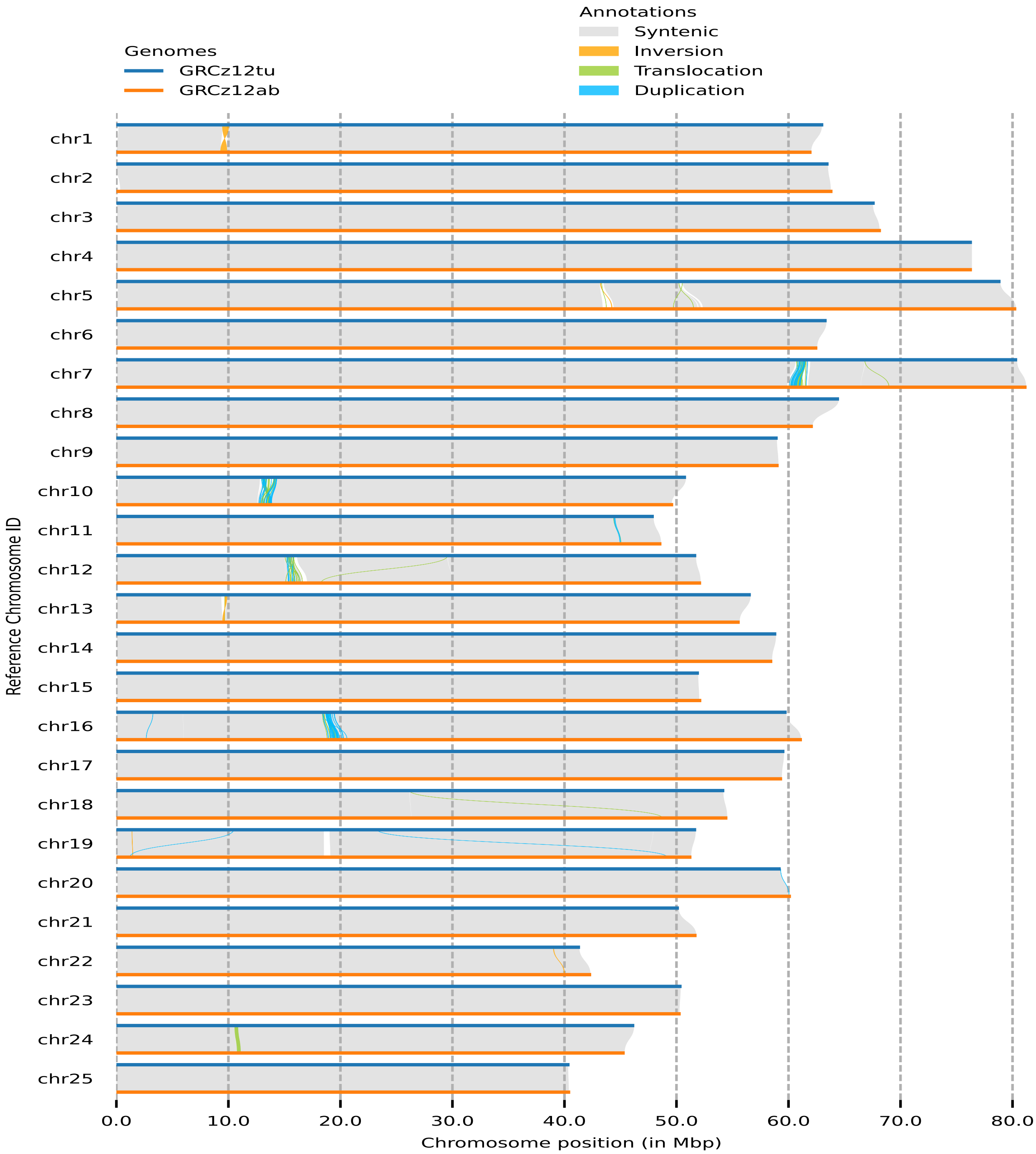

**Supplementary Fig 17:** Syntenic analysis and comparison between the GRCz12tu and GRCz12ab genome assemblies. The annotation differences such as duplications, translocation, and inversion only imply the segmental differences between the two genome assemblies.

#### References

1 Rautiainen, M. & Marschall, T. GraphAligner: rapid and versatile sequence-to-graph alignment. *Genome Biology* **21** (2020). https://doi.org/ARTN 25310.1186/s13059-020-02157-2

2 Rhie, A. *et al.* The complete sequence of a human Y chromosome. *Nature* **621** (2023). https://doi.org/10.1038/s41586-023-06457-y

3 Poplin, R. *et al.* A universal SNP and small-indel variant caller using deep neural networks. *Nature Biotechnology* **36**, 983-+ (2018). https://doi.org/10.1038/nbt.4235

4 Danecek, P. *et al.* Twelve years of SAMtools and BCFtools. *Gigascience* **10** (2021). https://doi.org/ARTN giab00810.1093/gigascience/giab008

5 Formenti, G. *et al.* Merfin: improved variant filtering, assembly evaluation and polishing via K-mer validation. *Nature Methods* **19**, 696-+ (2022). https://doi.org/10.1038/s41592-022-01445-y

6 Rhie, A., Walenz, B. P., Koren, S. & Phillippy, A. M. Merqury: reference-free quality, completeness, and phasing assessment for genome assemblies. *Genome Biology* **21** (2020). https://doi.org/ARTN 24510.1186/s13059-020-02134-9

7 Jain, C., Rhie, A., Hansen, N. F., Koren, S. & Phillippy, A. M. Long-read mapping to repetitive reference sequences using Winnowmap2. *Nat Methods* **19**, 705-710 (2022). https://doi.org/10.1038/s41592-022-01457-8

8 Li, H. Minimap2: pairwise alignment for nucleotide sequences. *Bioinformatics* **34**, 3094-3100 (2018). https://doi.org/10.1093/bioinformatics/bty191

9 Li, H. *et al.* The Sequence Alignment/Map format and SAMtools. *Bioinformatics* **25**, 2078-2079 (2009). https://doi.org/10.1093/bioinformatics/btp352

10 Smolka, M. *et al.* Detection of mosaic and population-level structural variants with Sniffles2 (vol 42, pg 1571, 2024). *Nature Biotechnology* **42**, 1616-1616 (2024). https://doi.org/10.1038/s41587-024-02141-2

11 Goel, M., Sun, H. Q., Jiao, W. B. & Schneeberger, K. SyRI: finding genomic rearrangements and local sequence differences from whole-genome assemblies. *Genome Biology* **20** (2019). https://doi.org/ARTN 27710.1186/s13059-019-1911-0

12 Benson, G. Tandem repeats finder: a program to analyze DNA sequences. *Nucleic Acids Research* **27**, 573-580 (1999). https://doi.org/DOI 10.1093/nar/27.2.573

13 Bedell, J. A., Korf, I. & Gish, W.:: a performance enhancement to RepeatMasker. *Bioinformatics* **16**, 1040-1041 (2000). https://doi.org/DOI 10.1093/bioinformatics/16.11.1040

14 Iseric, H., Alkan, C., Hach, F. & Numanagic, I. Fast characterization of segmental duplication structure in multiple genome assemblies. *Algorithm Mol Biol* **17** (2022). https://doi.org/ARTN 410.1186/s13015-022-00210-2

15 Lin, Y. Z. *et al.* quarTeT: a telomere-to-telomere toolkit for gap-free genome assembly and centromeric repeat identification. *Hortic Res-England* **10** (2023). https://doi.org/ARTN uhad12710.1093/hr/uhad127

16 Vollger, M. R., Kerpedjiev, P., Phillippy, A. M. & Eichler, E. E. StainedGlass: interactive visualization of massive tandem repeat structures with identity heatmaps. *Bioinformatics* **38**, 2049-2051 (2022). https://doi.org/10.1093/bioinformatics/btac018

17 Quinlan, A. R. & Hall, I. M. BEDTools: a flexible suite of utilities for comparing genomic features. *Bioinformatics* **26**, 841-842 (2010). https://doi.org/10.1093/bioinformatics/btq033

18 Ou, S. J. *et al.* Benchmarking transposable element annotation methods for creation of a streamlined, comprehensive pipeline (vol 20, 275, 2019). *Genome Biology* **23** (2022). https://doi.org/ARTN 7610.1186/s13059-022-02645-7

19 Porubsky, D. SVbyEye: A visual tool to characterize structural variation among whole-genome assemblies. (2024). https://doi.org/https://doi.org/10.1101/2024.09.11.612418

20 Rautiainen, M. *et al.* Telomere-to-telomere assembly of diploid chromosomes with Verkko. *Nat Biotechnol* (2023). https://doi.org/10.1038/s41587-023-01662-6

21 Hicktey, G. *et al.* Pangenome graph construction from genome alignments with Minigraph-Cactus. *Nature Biotechnology* **42** (2024). https://doi.org/10.1038/s41587-023-01793-w

22 Guarracino, A., Heumos, S., Nahnsen, S., Prins, P. & Garrison, E. ODGI: understanding pangenome graphs. *Bioinformatics* **38**, 3319-3326 (2022). https://doi.org/10.1093/bioinformatics/btac308
